# Deep models of protein evolution in time generate realistic evolutionary trajectories and functional proteins

**DOI:** 10.64898/2026.02.19.706898

**Authors:** Antoine Koehl, Sebastian Prillo, Matthew Liu, Junhao Xiong, Lillian Weng, David F. Savage, Yun S. Song

## Abstract

Models of protein evolution are foundational to biology, underpinning essential techniques such as phylogenetic tree inference, ancestral sequence reconstruction, multiple sequence alignment, variant effect prediction, and protein design. Historically, for computational tractability, these models have relied on the simplifying – but biologically unrealistic – assumption that sites in a given protein evolve independently of each other. A crucial test of any evolutionary model is its ability to simulate realistic evolutionary trajectories, but the independent-sites assumption leads to simulations that poorly reflect the complexity of natural protein evolution. Here we introduce PEINT (Protein Evolution IN Time), a flexible and generalizable deep learning framework for modeling how the entire protein sequence evolves over time while incorporating complex interactions between sites. This framework enables learning realistic patterns of constrained evolutionary transitions directly from millions of protein sequences spanning diverse fold families. Furthermore, unlike classical models that require pre-aligned sequences, PEINT learns indel dynamics directly from raw, unaligned sequences, thereby eliminating potential biases from alignment errors that can lead to incorrect inference of evolutionary patterns. By capturing higher-order epistatic interactions and modeling insertion-deletion processes that classical models typically ignore, PEINT accurately reproduces key signatures of natural evolution, including conservation patterns and family-specific dynamics. When simulating evolution along phylogenetic trees, PEINT generates highly novel sequences that preserve protein function, which we validate through experimental characterization of simulated carbonic anhydrase variants that retain enzymatic activity. PEINT thus enables realistic simulation of protein evolution that explores new sequence space while respecting structural and functional constraints. This evolution-informed generative modeling framework offers a powerful new tool for advancing both phylogenetic inference and protein engineering.

## Introduction

Models of protein evolution describe how protein sequences evolve over time under mutation and various selective pressures. These models are at the core of most applications in evolutionary biology, and act as the engine that powers a variety of phylogenetic methods, including phylogenetic tree inference [1–3], ancestral sequence reconstruction [4–6], multiple sequence alignment [7], variant effect prediction [8], and protein design [9], among others. Classical models of protein evolution date back to Margaret Dayhoff [10–12], and further refinements of her seminal work have proceeded in lock step with ever increasing data in sequencing databases [13–15]. These methods typically use continuous-time Markov chains (CTMCs) to model protein evolution. A CTMC is parametrized by a rate matrix (frequently denoted by *Q*) that describes the relative mutation rates and transition probabilities of different amino acids as a function of evolutionary time. Historically, efforts in the field have focused on estimating better rate matrices (Figure 1a). To date, the most widely used models of protein evolution are the WAG [14] and LG models [15], both parametrized by a rate matrix Q that is learned using maximum likelihood methods. The LG model generalizes WAG by adding the notion of site-rate variation, in which different sites are allowed to evolve with different overall rates, providing a means to deal with different patterns of conservation across sequences.

**Figure 1:**
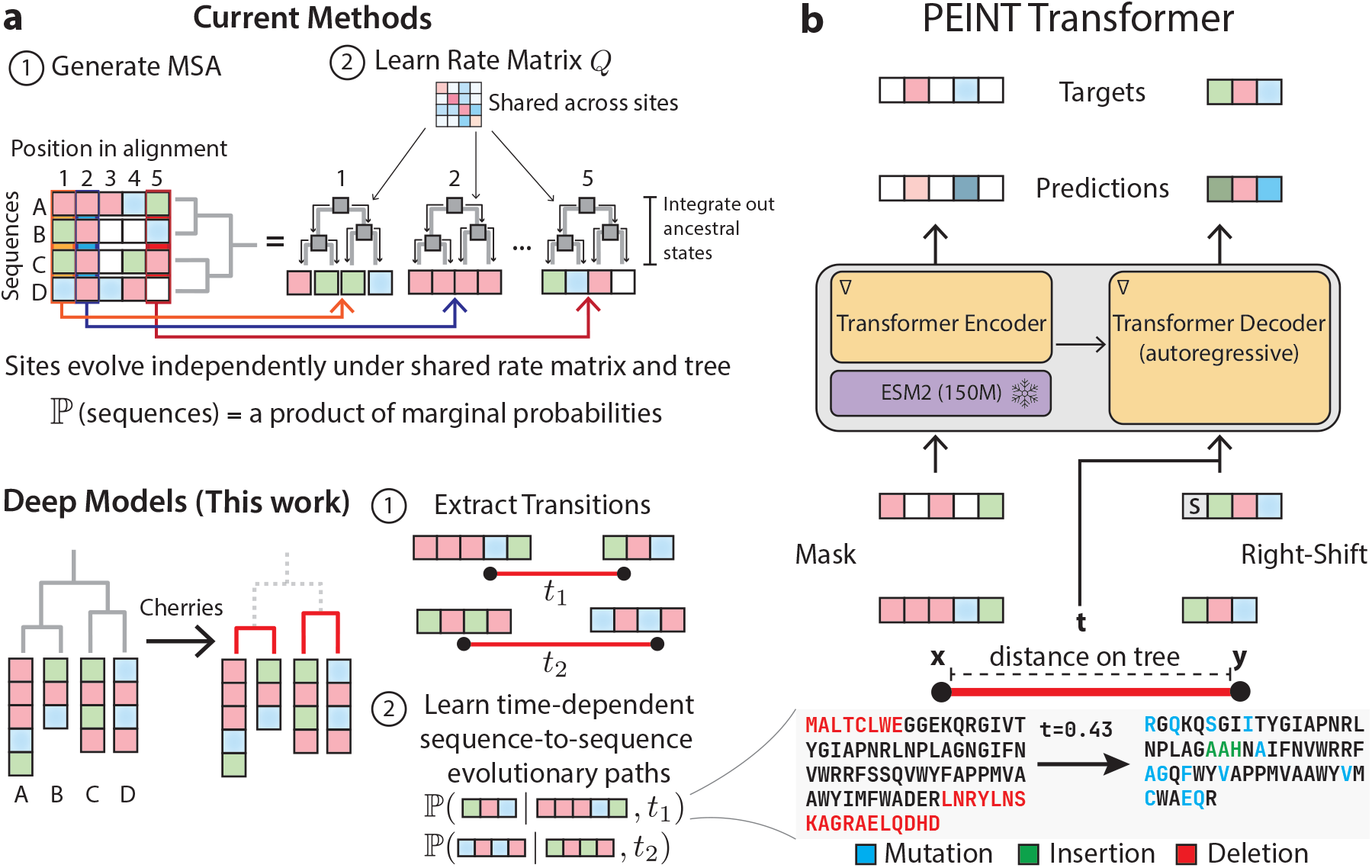
Learning models of protein evolution. Classically, models of protein evolution learn the parameters for a shared rate matrix that describes how each site in a protein’s multiple sequence alignment evolves over time (**a**, top). The problem of learning from evolution can be recast as learning transitions between full sequences in the cherries of the tree (**a**, bottom). Our PEINT transformer (**b**) learns to model evolutionary transitions in these cherries, accounting for mutations, insertions and deletions.

While highly successful, these classical models come with two important shortcomings: they require sequences to be pre-aligned with gap characters (‘–’) to ensure the same length, and assume that each site evolves independently of other sites. Although alignment errors can lead to incorrect inference of evolutionary patterns and the independent-sites assumption is inherently flawed, these simplifications have been standard in phylogenetics due to computational tractability. If a model of protein evolution is thought of as a translation between ‘past’ and ‘future’ versions of a protein, then the independent-sites assumption is akin to translating a sentence between languages one word at a time, independently of the other words in the sentence. Models that can move beyond this independent-sites assumption promise to improve performance on many downstream tasks.

In a different vein, recent applications of deep learning have made great strides in probabilistic models of protein sequences [16–19]. Though these models leverage deep learning architectures that are particularly adept at modeling interactions between sites, they cannot inherently describe how one protein evolves into another over time while experiencing intricate structural and functional constraints or adapting to new functions. These models implicitly treat sequences as being independent and identically distributed (i.i.d.); this assumption greatly simplifies the modeling task, but it is also a source of bias [20, 21]. The key issue is that the i.i.d. assumption fundamentally ignores the evolutionary history (or relatedness) of proteins and makes such models unable to reason about how proteins evolve over time. Empirically, this manifests as phylogenetic bias towards the most highly represented organisms in sequence databases [20, 21], which is typically addressed in a heuristic manner by prior clustering of the databases to remove sequences that are too similar to one another. Although this approach to evolutionary de-duplication is successful in mitigating issues that come from biased sampling of sequences across the tree of life, it discards local variation in sequences that is known to be important for specialized function in protein families that share a fold [22].

Here, we demonstrate how to achieve the best of both worlds: by integrating principled evolutionary reasoning with modern protein language models, we introduce a new framework for learning models of protein evolution in time that is alignment-free and accounts for higher-order interactions. This enables us to make use of epistatic signals within sequences to build more sophisticated models of variation between sequences. Importantly, we show that our approach can make use of evolutionary data at scale, leading to models that show significantly improved performance at modeling both retrospective and prospective evolution. We anticipate that further work along these lines will lead to novel insights about the evolution that has led to our current protein universe, as well as models that can more accurately predict forward evolution of proteins involved in disease processes.

## Results

### The PEINT framework for modeling sequence evolution

The objective of evolutionary modeling is to estimate a likelihood over a set of sequences that are related under a phylogenetic tree (for a more thorough treatment, see Supplementary Text in Supplementary Materials). The difficulty arises from the fact that we typically only have access to the leaves of the tree (extant sequences) and ancestral sequences must be integrated out. This computation is only tractable for independent-sites models [23], which, by definition, cannot accurately model higher-order epistatic interaction patterns that are known to be essential for protein structure and function. Recent work has shown that one can instead ignore the ancestral states completely, by using a composite likelihood over cherries, and learn models of protein evolution that move beyond the independent-sites assumption [24]. The composite likelihood approach recasts the problem of learning a complicated joint probability over many sequences to one of learning many conditional probability paths between pairs of sequences – termed “cherries” – on a tree. Intuitively, this can be thought of as decomposing a larger tree into many smaller trivial trees – “sticks” – in which both the root and leaf are observed, assuming time-reversibility. This allows us to train a model using a composite likelihood loss:

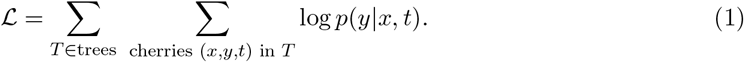

In (1), a triple (*x, y, t*) makes up a “cherry” and *p*(*y*|*x, t*) quantifies the conditional probability of observing a sequence *y* given a starting sequence *x* and the time t on the tree that separates them. The composite likelihood formulation has a key advantage: in sidestepping the ancestral state integration step, evolutionary transitions are no longer constrained to occur between aligned positions in a multiple sequence alignment. This has two important consequences. First, it allows for the use of arbitrary model classes, including deep neural networks, to learn directly from evolutionary transitions across entire sequences in one go. Secondly, it removes the constraints of relying on a pre-computed alignment, which may contain errors, and in doing so enables models to more natively learn from insertion and deletion (indel) processes within protein families. Despite their outsized importance in evolution [25], indels have traditionally been difficult to incorporate into classical models of evolution.

Here, we present the PEINT (Protein Evolution IN Time) framework, which, to our knowledge, represents the first implementation of a deep neural network to learn explicit models of protein evolution over time (Figure 1b). The PEINT model uses a coupled encoder-decoder transformer – this architecture is the standard in the field of machine translation [26] – and is aptly suited to learning under a composite likelihood framework which recasts the fitting of protein evolutionary models into a direct sequence-to-sequence setup [27]. Concretely, the transformer parametrizes the transition probability in a single cherry as *p*(*y*|*x, t*) = Π_*i*2sites(*y*)_ *p*(*y*_*i*_|*y*_1:*i*−1_, *x, t*) – that is, for each site i in sequence *y*, estimating its likelihood given the conditioning sequence *x*, the branch length t, and the already generated positions in *y* ({1,…, *i* − 1}). In our initial experiments, we found that overall model performance was correlated with the strength of the encoder module, which is designed to learn rich, structural representations of the input sequence x [28, 29]. We therefore reasoned that providing pretrained general purpose protein sequence representations from a state-of-the-art encoder protein language model (e.g., ESM2 [17]) as initial inputs to our network would enhance its overall performance. However, we note that as a general framework, PEINT is agnostic to the particular choice of encoder model, allowing for modular training of new models of protein evolution on any arbitrary pretrained BERT-style model [30]. Our final model, with ∼57M trainable parameters, builds a further 5 encoder and decoder layers on top of fixed representations from the 150M-parameter ESM2 model. The PEINT model was trained on a composite task of masked language modeling on the input sequence (*x*), and autoregressive (“next token prediction”) generation on the transition sequence (*y*).

### Data for learning evolutionary models using PEINT

To learn from evolutionary transitions within well-known protein folds, we turned to the TrRosetta dataset [31] – a set of ∼15,000 multiple sequence alignments (MSAs) that come from a sequence clustering of the PDB (Supplementary Figure S1a). We used these MSAs to infer evolutionary trees [3], and extracted evolutionary transitions (cherries) from these trees. To train and evaluate classical phylogenetic models, we retained the initial alignment in our transitions. For the PEINT model, we re-extracted the original sequences from UniProt and removed all gap symbols introduced by alignments, thus allowing the model to learn from native sequences.

For evaluation purposes, we used two levels of dataset splitting. In the first, we held out the sequences from ∼500 trees to evaluate generalization outside of the training domain (test families). For the remaining ∼14,500 families, we performed an in-family holdout split. This split involves identifying an early branch within the tree whose removal would yield two balanced subtrees. These subtrees show similar distributions of transition times, but are the result of independent evolutionary trajectories, and therefore contain sequences with low identity to one another (Supplementary Figure S1c). We held out one of the subtrees and used the other to train the model. These two splits enable us to assess generalization at different levels – to new sequences within protein families that we train on (in-family holdout) as well as new sequences in previously unseen protein families (test families). In sum, our dataset contains ∼14,500 training families with 512 cherries per family, totaling ∼15 million transitions. We used a conventional notion of time that is inherited from classical evolutionary models – rates are normalized such that one unit of time corresponds to there being one mutation per site in expectation (Supplementary Figure S1b). This dataset has a median transition t = 0.38 (Supplementary Figure S1d) and the pair of proteins in a cherry differ on average at ∼35% of sites – offering a rich source of information to learn about evolutionary forces in proteins. Models typically converged in a matter of 1-2 days on 2× A100 GPUs.

### Evaluating evolutionary transitions along single branches

A model of protein evolution should be able to reason over various combination of its inputs – that is, a pair of sequences (*x, y*) and the branch length that separates them (*t*). By varying which inputs were provided to PEINT, we assayed its ability to reconcile how evolutionary time is related to mutational burden across protein families.

First, we used the composite likelihood function on held-out transitions from real data. In this test, each model is provided the same set of evaluation cherries, and must assign a likelihood to these observed data. To build intuition for this task, we can compare with the uniform random guess model – a trivial model of evolution which assumes that each site in a protein will randomly mutate to any of the 20 amino acids across all timescales. This model would assign a uniform data likelihood of 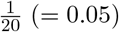 across all times (Figure 2a, dotted gray line). A more sophisticated model should be able to reason about the kinds of mutations that are most likely to occur (or not occur) over various evolutionary distances. To enable a direct comparison between PEINT, which operates outside the context of an alignment, and classical, alignment-bound models, we first calculate per-site transition likelihoods ℙ(*y*_*i*_|*y*_1:*i*−1_, *x, t*) for all models, but only take mean likelihood over positions that are present in the core alignment that the classical models of evolution are trained on (see Materials and Methods). We further stratified model performance as a function of the time separating the sequences in the test cherries. Among classical models, the LG model [15] shows better performance relative to the WAG model [14], as it explicitly models site-rate variation and can better account for patterns of conservation across sites in protein families (Figure 2a). The PEINT transformer shows superlative performance across the board (Figure 2a). Most importantly, this improvement persists even on transitions from trees held out during training (Figure 2a, right), highlighting PEINT’s ability to generalize evolutionary insights across protein families. We hypothesize that this increased capability comes from the use of a broadly pretrained protein language model (ESM2) [17] that helps to develop rich representations of protein sequences in the encoder part of the PEINT model [32]. Overall, PEINT’s unprecedented ability to model evolutionary transitions shows the promise of combining deep neural networks with the composite likelihood objective to learn from patterns of evolution across the proteome.

**Figure 2:**
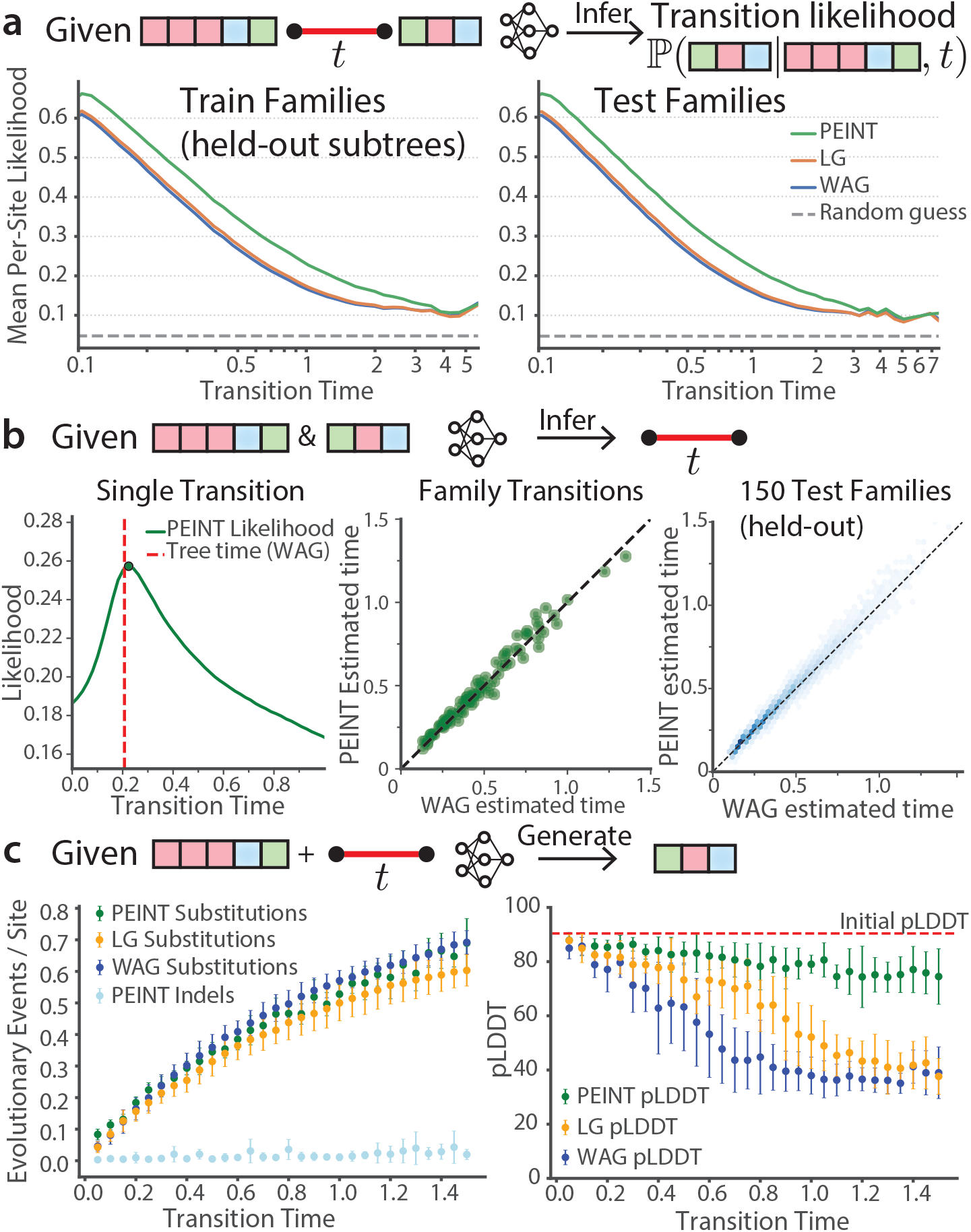
PEINT learns characteristics of evolution along single branches. Models of protein evolution must learn to relate sequences and evolutionary times. **a**) Given held out evolutionary transitions from subtree splits (left) and entirely held out trees (right), PEINT significantly outperforms classical models (WAG, LG) in terms of data likelihood over all evolutionary times. **b**) Given a pair of sequences *x* and *y*, PEINT can estimate the time separating them by maximizing *p*(*y*|*x, t*) + *p*(*x*|*y, t*) over *t* (left panel). This procedure can be applied at scale to pairs of sequences in a single tree (middle), as well as across 150 held-out families (right, Pearson’s *r* = 0.95). **c**) Given a starting sequence and time, PEINT generates novel sequences with defined mutational loads that mirror those of classical models (left). Compared to independent-sites models, PEINT generates structurally coherent sequences (right).

### PEINT can infer branch lengths

Intuitively, a model of protein evolution should learn a notion of the evolutionary time that separates sequences based on the number and types of mutations that occur between them. Classically, one can obtain a maximum likelihood estimate of the evolutionary time separating a pair of aligned sequences *x, y* in an MSA using classical models by maximizing *p*(*y*|*x, t*). This information is the basis for neighbor joining tree reconstructions algorithms [33, 34]. Similarly, one can use PEINT to estimate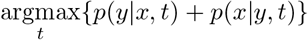 – that is, the time that maximizes the transition probability between a pair of observed sequences (in both directions), even in the absence of an explicit alignment. Empirically we found that PEINT generates concave likelihood curves as a function of time, so we used gradient ascent on the likelihood to solve this optimization problem (Figure 2b, left). Importantly, this is done in the absence of an alignment, thus mitigating the potential for alignment errors to lead to incorrect or biased estimates of divergence times. We carried out this procedure on cherries from held-out families – for a specific held-out family (Figure 2b, middle) and also across 150 randomly selected held-out families (Figure 2b, right) – and observed strong correlations between PEINT’s time estimates and those from classical models (Pearson’s r = 0.95). This strong correlation suggests that PEINT is able to provide consistent estimates of evolutionary distances within the time framework in which it is trained, even in trees it has not seen. As PEINT estimates are obtained from unaligned sequences, our results suggest the potential of deep models of protein evolution to more accurately reconstruct phylogenies that reflect the full spectrum of mutational processes – mutations, insertions and deletions.

### PEINT generates novel sequences with defined mutational loads while retaining high structural confidence scores

Models of protein evolution are fundamentally generative in that they can be used to propose a distribution over sequences given a starting sequence and a branch length (i.e., the evolutionary time separating those sequences). For classical models, this is simple and involves independently sampling from the time-specific transition matrix (obtained by exponentiating a time-scaled rate matrix) for each position in the protein. On the other hand, generating high-quality, coherent outputs from autoregressive models entails many technical considerations and remains an active area of research. In addition to well-known failure modes of autoregressive models trained via likelihood optimization (e.g., repetitive sequences, bland or nonsensical outputs), we wanted to ensure that the outputs of our model would not be overly restricted by the starting sequence. We experimented with a variety of sampling schemes, including beam search, *top-k* sampling and nucleus sampling [35], but found that these led to outputs with significantly less diversity than what one would expect in natural sequence evolution. Instead, we found that simply adding a special token (<monospace>eos</monospace>), signifying the end of the protein, to the end of our inputs at training time and decoding until the model outputs the eos token without further modification of the output probabilities was the most performant sampling scheme. Since PEINT is trained on full length, unaligned sequences, it can generate new sequences with novel insertions and deletions while maintaining the core regions related to central function.

We compared the generative capabilities of the classical models WAG and LG, as well as PEINT, by fixing a starting sequence (*x*) and decoding a distribution of sequences with progressively larger branch lengths (longer *t*) (Figure 2c). We evaluated these sequences for their mutational load, as well as the structural confidence score, pLDDT, assigned to them using OmegaFold [36], a state-of-the-art structure prediction model that uses single sequences as input. All model classes showed similar behavior in that the average mutational load increased monotonically with evolutionary time and generated comparable numbers of mutations across the evaluated time range (Figure 2c, left).

However, we found that the quality of generated sequences differed greatly, even controlling for numbers of mutations, with PEINT generating consistently high quality sequences even at high mutational loads (Figure 2c, right). On average, PEINT-generated sequences show high folding confidence (pLDDT scores above 80) across more than one full time unit, at which point, roughly half of the starting protein had been mutated. Both the WAG and LG models, on the other hand, show a sharp drop in sequence quality on this longer timescale, highlighting the limits of independent-sites mutational models. We surmise that the ability of a deep learning model, such as PEINT, to capture epistatic interactions allows it to generate compensatory mutations that account for already mutated positions and thus maintain functionally important contacts. This is a crucial aspect that is novel to deep models of evolution, and prompted further investigation into PEINT’s ability to simulate evolution within protein families.

### PEINT simulates realistic evolution on a tree

Using phylogenetic trees to reconstruct evolutionary trajectories of genes and organisms is the central method that enables comparative genomics. Since ground truth data are generally lacking, phylogenetic inference model development and benchmarking has historically relied on simulated data of evolution on fixed tree structures [37]. Unfortunately, the ability to realistically simulate protein evolution has been lacking, and existing methods, based on classical, independent-sites models, are known to lead to unrealistic sequences that are easily distinguished from naturally evolved sequences [38]. The constraints of classical model-based simulators manifest in overly smooth loss landscapes relative to real evolutionary data [39], leading to large generalization gaps [40]. Furthermore, classical simulators struggle to generate realistic insertion and deletion events [41], and in many cases rely on empirical observations of common insertions that must be extracted for each alignment [42].

Encouraged by PEINT’s ability to generate high-quality distributions of sequences, including indels, along single branches (Figure 2c), we reasoned that it could be extended to simulate evolution within protein families along entire phylogenetic trees. Simulating evolution down a tree entails successive rounds of generating child sequences from parent sequences, with each new sequence in turn serving as the parent for the next round. This task came with the uncertainty as to whether PEINT could maintain structurally coherent sequences through these successive rounds of generation, which drive the model farther from the extant sequences on which it was trained. Starting from ∼550 empirical tree/MSA pairs from the fully held-out families, we first selected the sequence with the median length to set as the new root of the tree. Since these trees are roughly balanced binary trees, we could then designate the subtree that does not contain the selected root as the evaluation set, as the sequences present in that subtree are the same distance from the new root in the original and re-rooted tree, and allow us to empirically account for the level of natural variation expected in real protein evolution on a per-family basis (Figure 3a). Each tree contained roughly 512 cherries, of which ∼256 cherries comprised an evaluation subtree. To simulate evolutionary trajectories down the tree, we applied the sampling scheme developed above. Starting from the root, we recursively generated the children of each node, eventually stopping at the leaves of the tree. For the classical model-based simulations, we used a state-of-the-art simulation engine, AliSim [43], driven by either the WAG or LG model. Overall, we found that the simulations carried out by PEINT and the classical models WAG and LG largely recapitulated first-order diversity measures across the ∼550 input trees, with highly congruent numbers of mutations generated across branches as the simulations progressed down the tree. (Supplementary Figure S2a,b). Furthermore, we found that all models, including PEINT, generated highly novel sequences. We performed comprehensive BLAST [44] searches between all simulated leaves in each family against the full nonredundant (nr) database, and tabulated the percent identity of the closest extant sequence. Across all simulated trees, we find that the sequences generated by WAG, LG and PEINT shared at most ∼50% sequence identity with any known sequence (Supplementary Figure S2c).

**Figure 3:**
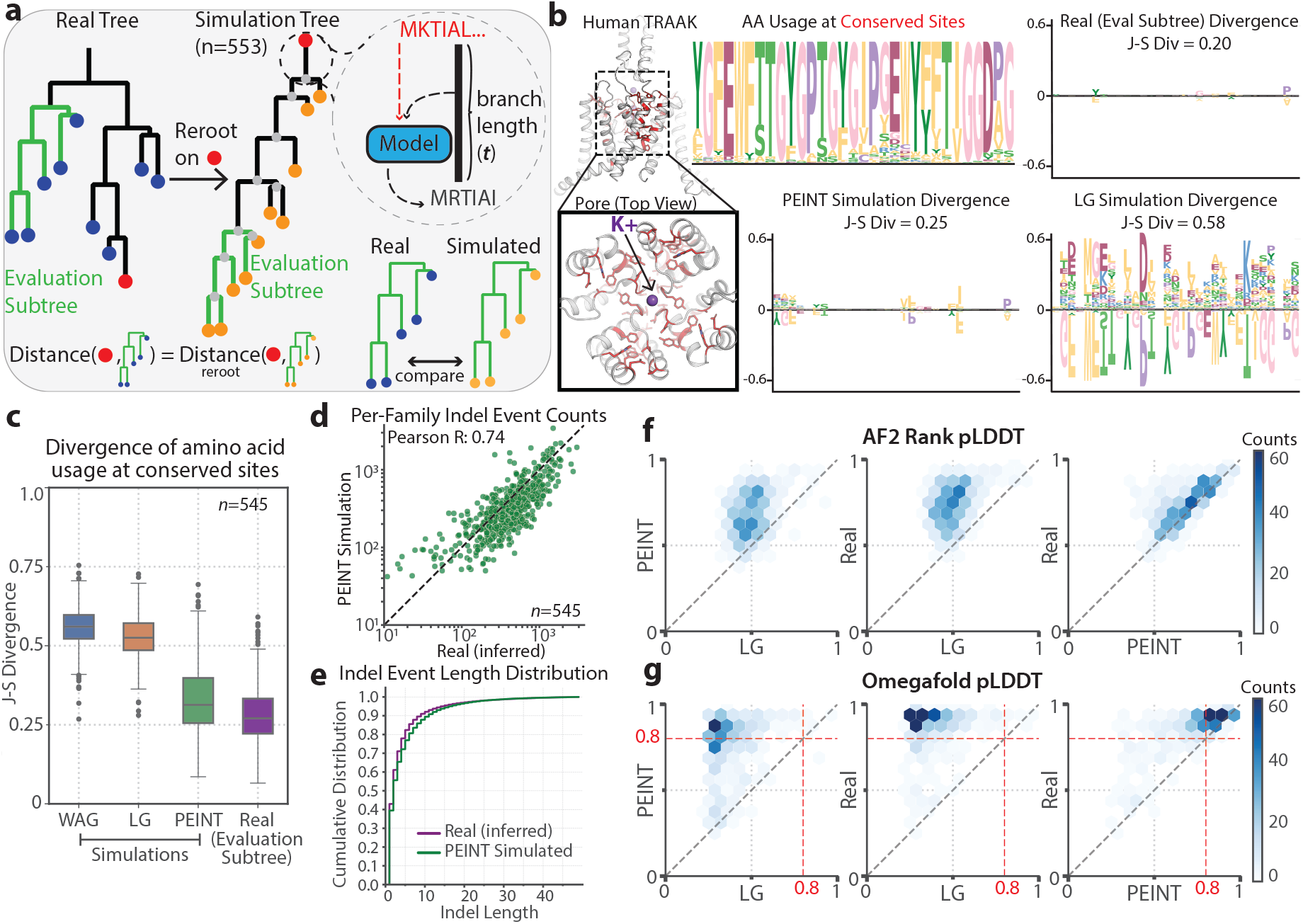
PEINT is a realistic simulator of protein evolution. **a)** Simulation setup – existing trees are re-rooted on a randomly chosen leaf sequence. From this root sequence, all internal and leaf nodes are progressively generated using classical (LG, WAG) or PEINT evolutionary models. To account for normal variation within each family, a subtree is retained for evaluation – its distance to the new root node remains unchanged. b) PEINT generates sequences that maintain amino acid usage at conserved sites, while simulations using classical models (LG) do not. Example shown for a single family (**b**), and aggregated across 545 simulation trees (**c**). PEINT faithfully recapitulates family specific insertion/deletion (indel) rates (**d**) and per-event insertion length statistics (**e**). When evaluated using AF2Rank (**f**) or OmegaFold (**g**), PEINT generated leaf sequences have pLDDTs that are almost identical to real sequences, and significantly higher than those generated using the best classical model (LG).

#### Amino acid patterns at conserved sites

We first examined the ability of our various simulators to faithfully recapitulate amino acid usage patterns at conserved sites in different proteins. For example, the mechanosensitive human TRAAK channel [45] shows a preponderance of conserved residues in the selectivity filter of its ion conducting pore, as well as pore-adjacent domains that are involved in regulation of channel currents [46] (Figure 3b). Starting from a multiple sequence alignment of TRAAK homologs, we identified the sites that show greater than 75% conservation. We then aligned the leaves from the evolutionary simulations carried out using LG and PEINT to this MSA, as well as the sequences from the held-out evaluation subtree, and quantified amino acid usage patterns at these positions. The expected level of divergence can be captured by examining the real sequences from the evaluation subtree, which show relatively little divergence relative to the other subtree, with a mean Jensen-Shannon (J-S) divergence across sites of ∼0.2 (Figure 3b). PEINT-simulated sequences show a similar level of divergence (J-S divergence of ∼0.25), mostly with subtle differences in non-pore residues that naturally show lower conservation in natural sequences. In contrast, the LG model – despite having access to conservation statistics through positional, family-specific site rates – struggles to maintain these functionally important sites across the tree, and generates sequences with highly dissimilar patterns of amino acid usage across all conserved positions, reflected in a much higher J-S divergence of ∼0.58. In general, mutations at highly conserved sites are not tolerated, and it is likely that this level of divergence would lead to wholly nonfunctional proteins. These results hold over the remaining 545 held-out simulation trees (Figure 3c): PEINT accurately recapitulates real patterns of amino acid usage at conserved sites, while both WAG and LG show dramatically divergent amino acid usage profiles at these highly conserved positions for each protein family.

#### Local structural motifs

A similar analysis can be carried out on the conservation of structural motifs in the form the FoldSeek 3Di states [47]. The FoldSeek 3Di alphabet reflects local structural environments that each amino acid in a protein will find itself in, providing a convenient string representations of three-dimensional structure. Across all simulations, we find that PEINT accurately maintains conserved structural motifs that define family folds, showing divergence that is on par with natural variation (Supplementary Figure S3a,b). On the other hand, both LG and WAG generate sequences with highly divergent 3Di profiles, suggesting that the mutations they introduce are likely to lead to disruption of three dimensional structure, and loss of fold identity.

#### Indel patterns

We next examined whether PEINT simulations faithfully recapitulated family-level insertion and deletion (indel) characteristics. Since each branch was simulated without constraint, PEINT can generate sequences of varying lengths by choosing to insert or delete residues along a parent-child pair. We compared indel statistics in the evaluation subtree of real and PEINT-simulated sequences. To infer these quantities in the real trees (for which we do not have access to internal nodes), we used Historian [48, 49], a state-of-the-art statistical alignment and phylogenetic reconstruction package. Even though these families were held out from the training set, we find that PEINT simulations accurately reflect family-level indel rates (Figure 3d), and that this correlation is only partially explained by sequence length (Supplementary Figure S3c,d), indicating that PEINT can match the characteristic indel frequencies of different protein families. Furthermore, we evaluated whether PEINT had learned a notion of characteristic indel lengths – that is, the number of amino acids inserted or deleted at each new event. Aggregated across the ∼540 tree simulations, we observe exceptional concordance between PEINT-generated and real indel lengths (Figure 3e).

### Structural confidence scores

Finally, we evaluated the simulated leaf sequences using structural metrics. For each simulated family, we randomly picked 30 leaf sequences for which to predict structures using AF2Rank [50] and OmegaFold [36] and noted the median model confidence (reported as pLDDT) for that family. AF2rank evaluates structure and sequence congruity by generating a predicted structure with AlphaFold2 [51] using a given structural template and a single sequence. By not providing homologous sequences, AF2Rank is able to remove biases from homology searches, which can either help or hinder predictions, and has been shown to evaluate the energetic favorability of a particular sequence adopting a given structure. On the other hand, OmegaFold does not use templates, and indirectly uses homology by predicting structures from single sequences using pretrained language model representations. Both metrics show that PEINT simulates evolution while maintaining structure, with pLDDTs that are distributionally like those of real sequences from the same families.

As shown in Figure 3f,g, even OmegaFold, which may struggle with sequences that show high divergence to the distribution of natural sequences it was trained on, nonetheless assigned high structural confidence scores to PEINT-simulated leaves, with pLDDT scores above 80 which is typically the chosen cutoff of highly confident structural predictions. On the other hand, sequences generated by using LG as a simulation engine show poor structural metrics, with median pLDDT scores around 50, which indicates poor probability of being folded.

To mitigate confounding effects of homology biases and the boost of a strong template, we employed AlphaFold2 [51] as an unbiased discriminator by directly providing the MSA calculated from the leaves of our simulations for structure prediction. We find that MSAs based on WAG and LG-driven simulations lead to poor structural confidence, whereas PEINT generates MSAs that are essentially indistinguishable from real for the purposes of structure prediction (Supplementary Figure S4).

Taken together, the results described above demonstrate that PEINT simulates evolution in a manner that is virtually indistinguishable from real evolution.

### PEINT generates functional enzymes with low sequence identity from known proteins

Carbonic anhydrases (CAs) are a ubiquitous family of enzymes that catalyze the hydration of carbon dioxide into aqueous bicarbonate. These enzymes balance an exceptional turnover rate (*k*_*cat*_) with poor affinity for substrate (*K*_*M*_) – carbon dioxide is a small, linear molecule with negligible dipole moment – and the majority of CAs have K_*M*_ values in the millimolar range, far above the normal level of partitioning of atmospheric CO_2_ into water (∼micromolar). We reasoned that carbonic anhydrase would present a stringent test of PEINT’s ability to realistically simulate evolution while preserving function across large sequence distances. To ensure the generalizability of these results, we selected a held-out tree, and simulated from its seed sequence – that of the β CA sequence from cultivated pea (*Pisum sativum*) (Figure 4a). While other *β* carbonic anhydrases were present in the training set, by design they show minimal sequence identity to this family, as they come from stringent clustering of the underlying fold space. We then applied our evolutionary simulation protocol on a pruned tree, yielding 24 designed leaves. All leaves showed exceptional designability metrics, with OmegaFold pLDDT values above 85. Furthermore, we note an absolute conservation of the catalytic site across all generated sequences, and structural predictions of these sequences in the presence of zinc confidently return the correct coordination geometry required for catalysis (Figure 4b). We compared PEINT-generated sequences to those from a parallel simulation in which the classical LG model was used as the evolutionary engine. While both simulations yielded sequences with similar accumulation of mutations relative to the root (Figure 4c, top panel), sequences generated using the LG model had notably worse structural confidence scores, with a median pLDDT of around 30. We also highlight how the majority of leaves generated using either model showed 40–50% sequence identity to any known gene in the NCBI nonredundant (nr) database, indicating how PEINT naturally explores new regions of sequence space in the process of simulating evolution down a tree. Based on these in-silico designability metrics, we randomly selected 20 leaves from the PEINT simulation tree for experimental characterization.

**Figure 4:**
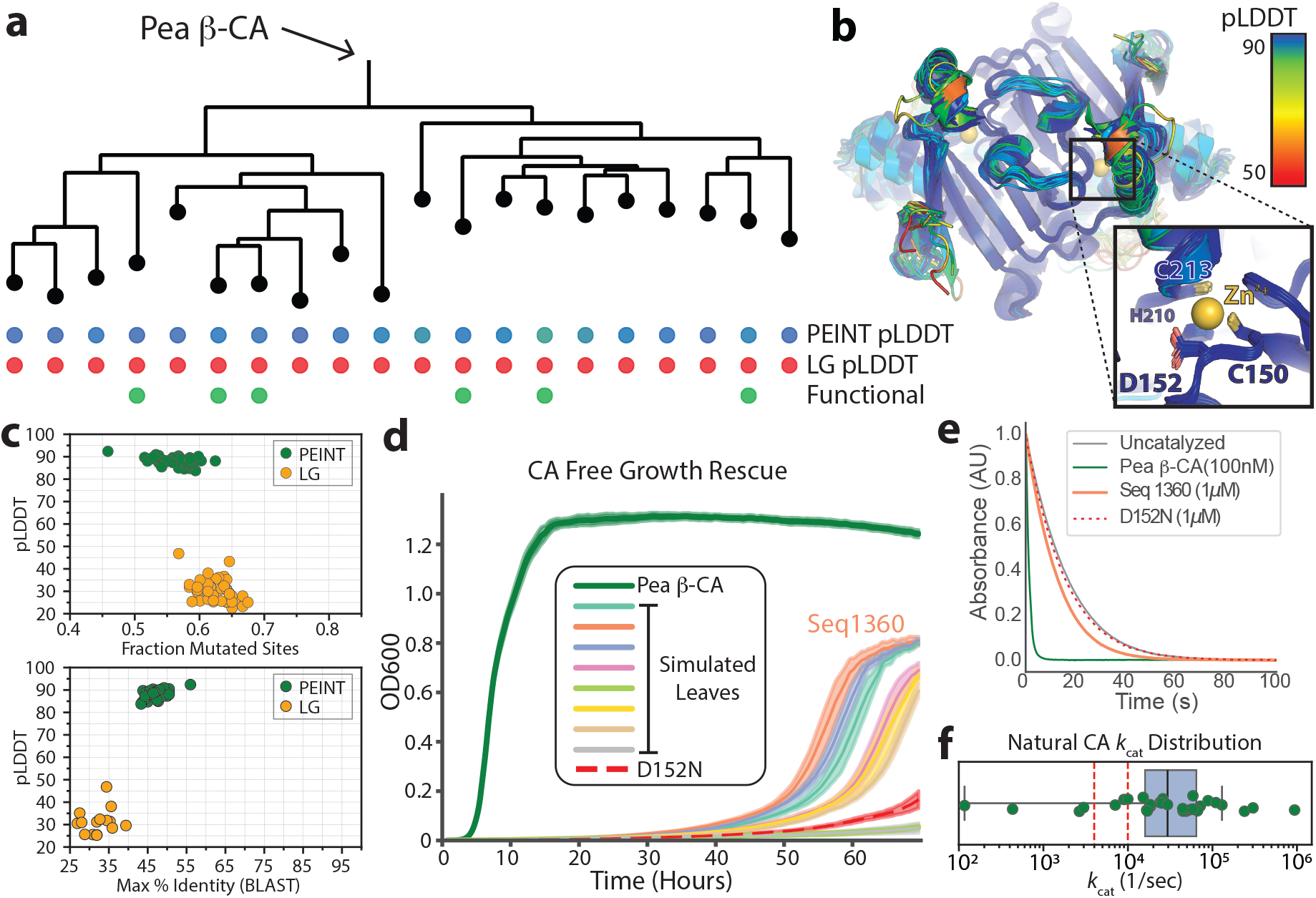
PEINT evolutionary trajectories maintain protein function at large sequence distances. **a**) Starting from the sequence of the Pisum sativum (Pea) *β*-carbonic anhydrase, PEINT and LG simulations were carried out on the same tree without filtering. **b**) Structural predictions of the PEINT generated leaves are shown, colored by pLDDT, and indicated below the tree in a. All sequences maintain the catalytic site (inset zoom) but show structural variation at neighboring sites that is predicted to maintain protein fold with high confidence. In comparison, the LG simulation leads to sequences with poor structural confidence metrics (pLDDT dots, **a**). **c**) Top - Both PEINT and LG-simulated sequences show similar numbers of mutations compared to the Pea *β*-CA, but LG generated sequences have significantly worse structural confidence (pLDDT) scores. Bottom - PEINT-generated sequences show low sequence identity to any extant sequence in BLAST databases, despite their high pLDDT. d) Growth over time in the CA-Free functional complementation assay. Six simulated leaves show growth between that of the pea *β*-CA and D152N mutation. **e**) In-vitro CO_2_ hydration shows a similar trend. **f**) Distribution of *k*_*cat*_ parameters for natural carbonic anhydrase enzymes span a large range. To roughly situate PEINT-simulated sequences, empirical *k*_*cat*_ values of D152N site mutations are shown (red lines).

As an initial assay to test the functionality of the simulated CA genes, we used a functional complementation assay in the Escherichia coli CAfree [52, 53] background. This engineered bacterial strain is deficient in endogenous carbonic anhydrase genes and cannot grow in low CO_2_ concentration in the absence of an exogenously provided functional CA gene. We transformed the 20 selected leaves, the root sequence [54] as a wild-type control, and a previously characterized mutant at a conserved aspartic acid residue, D152N (D162N in our construct). Mutations at site 152 have been previously shown to impair function by limiting the rate of the proton transfer step during catalysis, yielding slower, but still marginally functional carbonic anhydrases [55–57]. At an intermediate, but not growth-permissive, CO_2_ concentration of 1%, we found that 6 of the 20 designs could rescue growth better than the D152N mutation (Figure 4d), though all intriguingly showed a long lag phase phenotype. We followed this growth-based functional complementation assay with an in-vitro characterization of CO_2_ hydration rates. We expressed and purified the wild type pea *β*-CA, the D152N mutant, as well as one of our growth-rescuing simulation endpoints, seq1360, and measured the rate of hydration using the pH indicator assay [58] at a saturating concentration of 17 mM CO_2_ (Supplementary Figure S5). This assay validated our results from functional complementation, with seq1360 showing a slower rate of hydration than the wild type enzyme (despite a higher concentration), but faster reaction kinetics than the D152N mutant, which is only marginally faster than the uncatalyzed reaction (Figure 4e).

Despite a historical notion that CA is essentially a perfect enzyme, carbonic anhydrases across the tree of life are not universally perfect in their rates [59], with many possessing more modest k_cat_ values in the ∼10^4^*s*^−1^ range (i.e., roughly one percent of the well-known human CAII enzyme) (Figure 4f). While we did not measure full kinetic curves, we can approximately place our functional PEINT-generated sequences relative to independent measurements of the kinetics of mutations at the conserved aspartic acid (D152) (Figure 4f, red dashed lines), which implies that our designed enzymes’ k_cat_ values should fall within the range that is naturally observed in different organisms. It is possible that, by training more broadly on evolutionary transitions across the tree of life, PEINT has learned patterns of evolution from organisms for which the rapid hydration of CO_2_ is not as imperative as it is in human and plant metabolism, and may predominantly generate enzymes with modest, but measurable activity.

### PEINT improves variant effect prediction (VEP)

Since the need to maintain function constrains protein evolution, extant protein families contain a partial record of millions of years of evolutionary trial and error selecting for a given function. Intuitively, a strong model of protein evolution will natively be able to learn from protein family data, and thus should be able to predict the effect of mutations in different protein families. Density models fit on evolutionarily related sequences, either for specific families [60, 61] or across the whole tree of life [62, 63] have shown to be strong *zero-shot* variant effect predictors, which means that they do not require access to assay-labeled training data in order to score variant effects. In particular, large-scale protein language models (pLMs) such as ESM2 have shown to be strong predictors of variant effects [17], and are now being scaled to billions of parameters. However, recent works also suggest that increasing scaling of pLMs do not necessarily translate to increased performance on VEP tasks [64, 65], potentially due to its likelihood being biased towards certain evolutionary neighborhoods [20], suggesting that better accounting for phylogenetic relationships might be a promising approach for improving the performances of these models for VEP.

Herein we explored ways to leverage the PEINT framework to predict the effect of substitutions in families from ProteinGym [66], a standardized benchmark for protein VEP. We utilized the MSAs provided by the ProteinGym benchmark for these families and followed the same procedure as for the TrRosetta dataset to infer evolutionary trees and extract evolutionary transitions from these MSAs. To assess whether PEINT can leverage evolutionary information specific to these families, we trained a smaller PEINT model with ∼23M trainable parameters on top of the 150M-parameter ESM2 model, using a collection of 1,000 transitions from each family in the ProteinGym substitution benchmark. The model typically converges in a few hours on a single A100 GPU.

Traditionally, when probabilistic models of proteins are used for VEP, a log-likelihood ratio between the mutant and wild-type sequence is used for scoring. These models provide either an exact or approximate likelihood estimate of sequences, which we will denote as *p*(*x*). Concretely, letting *x*^*mut*^ denote the mutant and *x*^*wt*^ the wild-type sequence. The score of the mutant is typically computed as 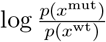, or, in the case of masked language models, as the conditional probability *p*(*x*_*M*_ |*x*_−*M*_) where *M* is the set of masked indices corresponding to the mutated positions. With an evolutionary model in hand, we can take a different approach to VEP. Following [67], we score mutant via 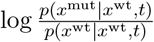, where *t* is a hyperparameter controlling that size of the evolutionary neighborhood around the wild-type. Note that within a given protein family, the denominator would be the same for all mutants and can therefore be ignored. We observed that for different protein families, the time which leads to the best Spearman correlation with the assay-labeled data differs, although varying the values of *t* within the typical ranges of the training distribution did not significantly change the overall results. Intuitively, this approach is appealing as it allows one to condition on the wild-type with varying degrees depending on the parameter *t* [67]. At *t* = 0, all mutants should have zero probabilities. As *t* increases from 0, mutant sequences will be assigned nonzero probabilities, and the extent to which these probabilities change depends on how likely the mutant could have arisen from the process of mutation and selection according to the model. Traditional scoring approaches with *p*(*x*) may be thought of as a particular instantiation of our approach with *t* = ∞, where the stationary distribution of the evolutionary model is used for scoring and the wild-type is ignored. While some approaches have been proposed to improve VEP with pLMs by accounting for wild-type preference [64] or injecting information from homologous sequences at inference time [65], our approach can be viewed as a complementary approach with the potential benefit of more fully leveraging the phylogenetic information from evolutionary data with a flexible model coupled with a principled training objective.

Each family in the ProteinGym substitution benchmark has been categorized by the assay type of the deep mutational scanning (DMS). Following the convention of the benchmark, we computed the Spearman correlation for each family, then examined the Spearman correlation aggregated over all families for each assay type as the primary metric. Using *t* = 1 as in Prillo et al. [67], we observed that the PEINT model trained on evolutionary transitions extracted from the MSAs of these families improves VEP performance over ESM2, which provided learned representation for the encoder of PEINT (Figure 5a). We also examined the results for mutants with varying distance from the wild-type sequences, and found that PEINT improves the performance of ESM2 at all mutational depths, particularly when the mutant sequences have more than two mutations from wild-type (Figure 5b). We hypothesized that this might be because PEINT models the likelihood of the entire sequence, which can better capture the joint distribution of multiple mutations than the conditional likelihoods from ESM2.

**Figure 5:**
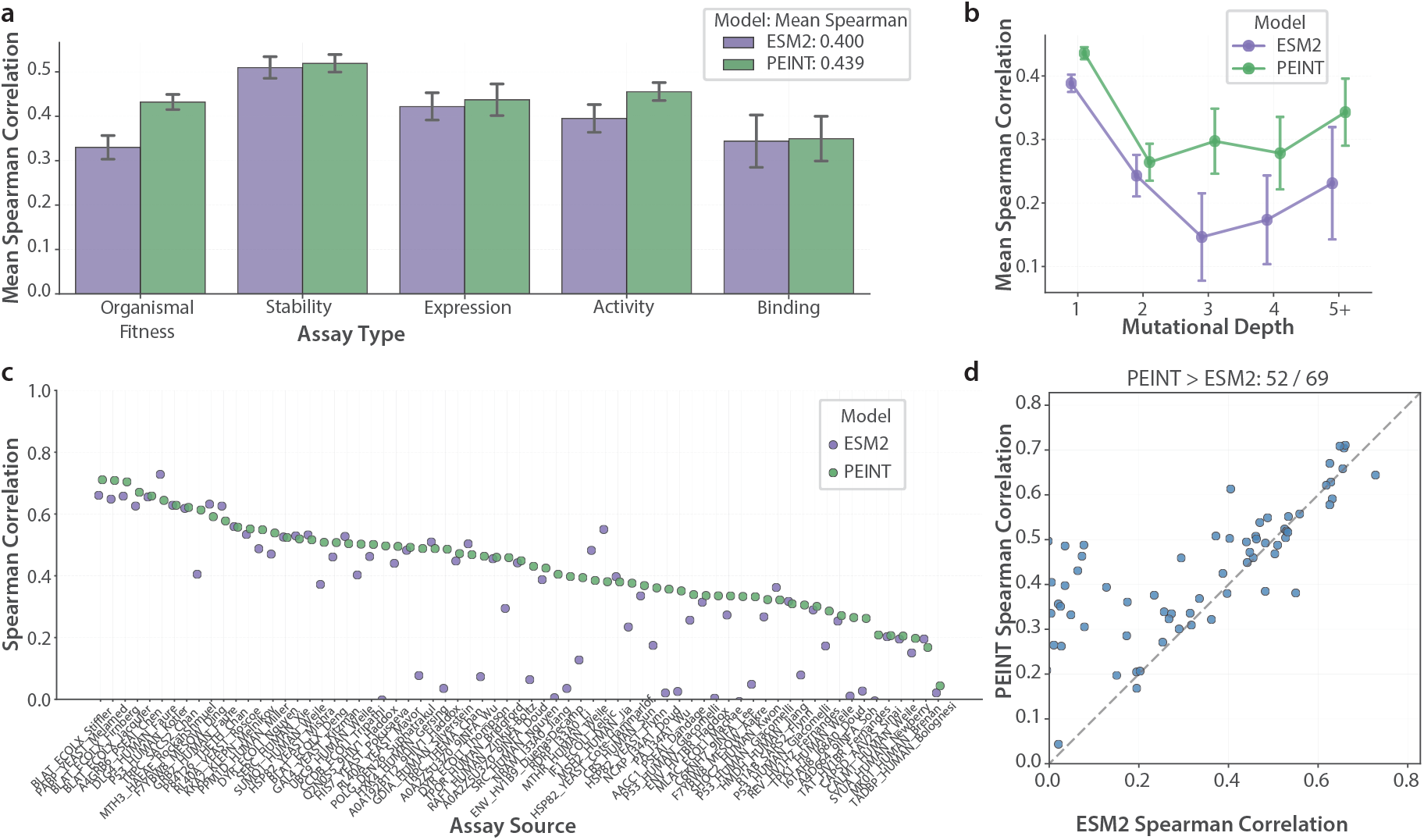
PEINT improves variant effect prediction. **a**) Mean Spearman correlation on 201 families in the ProteinGym substitution benchmark [66]. The families are stratified by assay type. The height of each bar represents the mean of the per-family Spearman correlations for each assay type. Error bars represent the standard deviation across proteins within each assay type. For each model, the average over the mean Spearman correlations for the five assay types is shown in the figure legend. “ESM2” is the 150M parameters ESM2 model, and “PEINT” is a PEINT model trained on transitions of the DMS families, using the frozen 150M parameters ESM2 in its encoder. **b**) Mean Spearman correlation stratified by mutational depths. **c**) Spearman correlation for each family in the ProteinGym substitution benchmark with the assay type “Organismal Fitness”, where the assays measure the extent to which mutations affect an organism’s growth rate. Each protein family is denoted by its identifier in the ProteinGym dataset. d) ESM2 vs. PEINT Spearman correlation for each family in the ProteinGym substitution benchmark with the assay type “Organismal Fitness”.

We found that PEINT offers the most significant performance gains on datasets where the assays measure the extent to which mutations affect an organism’s growth rate (Figure 5a). The growth rate is a more holistic proxy for the overall fitness constraints imposed on an organism compared to other molecular phenotypes (e.g., binding and stability), hence it stands to reason that a model that is better able to extract information from evolutionarily related sequences would be a better zero-shot variant effect predictor. Intriguingly, we saw that PEINT was able to rescue performance in many families where ESM2 struggled to generate reasonable predictions (Spearman *ρ* < 0.2) (Figure 5c,d).

Since the PEINT framework refines the learned representation of ESM2 in its encoder, it can in principle leverage more performative encoders to achieve better performance. To test this hypothesis, we trained another PEINT model on top of the 650M-parameter ESM2 model, which has better zero-shot performance than the 150M-parameter ESM2 model, and found that the resulting PEINT model improves upon its VEP performance even further (Supplementary Figure S6). The PEINT framework is highly flexible in its model architecture and can in principle also be applied to other pLMs. In this light, one can view PEINT as a general and principled framework to align pre-trained pLMs with evolutionary information and thereby improve their performance on VEP.

### Mechanistic interpretation of PEINT

Prior studies in mechanistic interpretability of protein language models have primarily focused on encoder-only (BERT) architectures, revealing that the masked language modeling objective naturally drives such models to learn rich sequence-structure relationships [29, 32, 68]. Since PEINT uses the more elaborate encoder-decoder architecture, we focused our attention on the decoder module, as well as inter-module communication. The ‘categorical Jacobian’ calculation, first introduced in Zhang et al. [32], systematically mutates each position in an input to a deep neural network and measures how these perturbations propagate through the model. Importantly, it provides a fully unsupervised means to evaluate the inner workings of models that operate on protein sequences. We first applied this technique to investigate PEINT’s Transformer decoder module. In particular, we wanted to ascertain whether PEINT was able to learn from and use epistatic interactions between sites in protein sequences to make more informed modeling choices at decoding time. Since the PEINT decoder works in an autoregressive manner, we systematically perturbed positions in the input to the decoder (i.e., the already-generated tokens) and measured their effect on downstream model outputs (Supplementary Figure S7a). This decoder categorical Jacobian [32] calculation reveals a highly structured pattern of inter-residue couplings that captures the topological structure of a protein’s contact map (Supplementary Figure S7b). Evaluated on a larger scale, we find that the most significant couplings learned by the PEINT decoder do correlate with structure, but more weakly than those learned by encoder-only models such as ESM2 (Supplementary Figure S7c). We hypothesize that functionally linked, but structurally dispersed, allosteric networks may be a large source of false positives when assayed against a static contact map. Indeed, prior work showed that the outputs of ‘mutant cycles’ carried out using autoregressive models are less attuned to direct structural contacts than masked language models [69].

We next investigated how the representations learned by PEINT’s encoder are shared with its decoder module. In the PEINT model for *p*(*y*|*x*, t), we systematically identified positions in the output y that were most affected by perturbations in the input sequence x (Supplementary Figure S7d). The cross-module categorical Jacobian couplings strongly resembled pairwise alignment trace matrices, suggesting that a large driver of PEINT’s ability to model evolution comes from its learned ability to align representations of the target sequence *y* with the source sequence *x* during decoding (Supplementary Figure S7e). Across ∼900 examples of curated alignments from the pFamA seed alignments[70], we find that PEINT encoder-decoder categorical Jacobian couplings natively recover ground truth alignments with similar accuracy as gold standard dynamic programming based alignment algorithms [71] (Supplementary Figure S7f), despite not being explicitly trained to do so. Taken together, these results suggest that the PEINT encoder and decoder modules serve fairly distinct purposes: the encoder builds a structural representation of the input sequence, while the decoder learns to align the target sequence to this representation while also learning patterns of evolution reflective of subfamily specialization.

## Discussion

Here we show, for the first time, how to use the composite likelihood objective over pairs of sequences to train a deep model of protein evolution with the PEINT Transformer. By moving beyond the independent-sites assumption that has stymied classical models for decades, PEINT demonstrates superlative performance across all facets of evolutionary modeling. The transformative potential of deep models of protein evolution is perhaps best exemplified by PEINT’s capacity for prospective evolutionary simulations. We demonstrate that PEINT naturally simulates evolutionary trajectories that explore new regions of sequence space while maintaining fold and function. The ability to generate realistic evolutionary trajectories at will has many implications for evaluating and improving phylogenetic inference methods, and synergizes particularly well with recent forays into using deep neural networks for this task [72]. Along similar lines, realistic simulations allow us to “replay the tape of life” under controlled conditions and can help generate testable hypotheses about the evolutionary forces that shape protein families [6, 73]. By innately capturing higher-order epistatic interactions and insertion-deletion processes that classical models ignore, deep evolutionary models promise to provide new avenues to advance our understanding of how proteins evolve in nature.

While classical models predominantly learn general trends of evolutionary drift, deep models of protein evolution can capture both general and family-specific selective pressures critical for maintaining function. This dual capability endows models such as PEINT with the ability to balance exploration and exploitation – as evidenced by maintaining function in evolutionary trajectories that explore new regions of sequence space that are substantially different from known proteins. This outcome naturally lends itself to protein engineering, especially within the machine learning-guided directed evolution framework [74]. In this context, the ability to generate diverse starting points enables design efforts that can more efficiently traverse a protein’s functional landscape. Current methods aiming to better align pLMs with variant effect data show mixed results. Simple model scaling yields diminishing returns and frequently leads to overfitting [20, 21], while protein-specific fine tuning can be challenging and unstable [62]. By conditioning on wild-type sequences and evolutionary time, PEINT provides an encoder-agnostic framework for aligning general protein language models with family-specific evolutionary information. The PEINT framework offers a principled alternative to prior approaches, with a lightweight training scheme that yields substantial performance gains in a matter of hours on a single GPU. These benefits are most pronounced when evaluating variants encompassing multiple mutations, as well as on assays representative of general protein function, and showcase how PEINT can tune broadly pretrained pLMs by grounding them in local evolutionary neighborhoods. Although we employed the ESM2 model in this work, we note that PEINT is a general framework that can utilize any pretrained BERT-style pLM as the encoder model.

In sum, our work lays the foundation for understanding and training deep models of protein evolution on sequence data at scale, and demonstrates their effectiveness across key tasks relevant to interpreting natural protein evolution. We anticipate that future work will refine this framework for specific applications of interest.

## Acknowledgments

We acknowledge members of the Song lab and the Savage lab for feedback on the manuscript, and Nitin Daniel and Andreas Martin for help with the stopped-flow system. We thank Martin Steinegger and Eli Levy Karin for helpful discussions on our work. This research is supported in part by NIH grants 5K99GM152766 and R35-GM134922, and by the UC National Laboratory Fees Research Program of the University of California Office of the President (UC AI Science at Scale Grant L26CR10102). David F. Savage is an investigator with the Howard Hughes Medical Institute.

## References

1. Minh, B. Q. et al. IQ-TREE 2: New Models and Efficient Methods for Phylogenetic Inference in the Genomic Era. Molecular Biology and Evolution 37, 1530–1534 (Feb. 2020).

2. Stamatakis, A. RAxML version 8: a tool for phylogenetic analysis and post-analysis of large phylogenies. Bioinformatics 30, 1312–1313 (Jan. 2014).

3. Price, M. N., Dehal, P. S. & Arkin, A. P. FastTree 2–approximately maximum-likelihood trees for large alignments. en. PLoS One 5, e9490 (Mar. 2010).

4. Yang, Z., Kumar, S. & Nei, M. A new method of inference of ancestral nucleotide and amino acid sequences. Genetics 141, 1641–1650 (1995).

5. Koshi, J. M. & Goldstein, R. A. Probabilistic reconstruction of ancestral protein sequences. Journal of Molecular Evolution 42, 313–320 (1996).

6. Starr, T. N., Picton, L. K. & Thornton, J. W. Alternative evolutionary histories in the sequence space of an ancient protein. Nature 549, 409–413 (2017).

7. Henikoff, S. & Henikoff, J. G. Amino acid substitution matrices from protein blocks. Proceedings of the National Academy of Sciences 89, 10915–10919 (1992).

8. Laine, E., Karami, Y. & Carbone, A. GEMME: a simple and fast global epistatic model predicting mutational effects. Molecular Biology and Evolution 36, 2604–2619 (2019).

9. Spence, M. A., Kaczmarski, J. A., Saunders, J. W. & Jackson, C. J. Ancestral sequence reconstruction for protein engineers. Current Opinion in Structural Biology 69, 131–141 (2021).

10. Eck, R. V. & Dayhoff, M. O. Evolution of the structure of ferredoxin based on living relics of primitive amino Acid sequences. en. Science 152, 363–366 (Apr. 1966).

11. Schwartz, R. M. & Dayhoff, M. O. Origins of prokaryotes, eukaryotes, mitochondria, and chloroplasts: A perspective is derived from protein and nucleic acid sequence data. en. Science 199, 395–403 (Jan. 1978).

12. Atlas of protein sequence & structure, suppl. No. 3 (ed Dayhoff, M. O.) (National Biomedical Research Foundation, Apr. 1979).

13. Jones, D. T., Taylor, W. R. & Thornton, J. M. The rapid generation of mutation data matrices from protein sequences. en. Comput. Appl. Biosci. 8, 275–282 (June 1992).

14. Whelan, S. & Goldman, N. A general empirical model of protein evolution derived from multiple protein families using a maximum-likelihood approach. en. Mol. Biol. Evol. 18, 691–699 (May 2001).

15. Le, S. Q. & Gascuel, O. An improved general amino acid replacement matrix. en. Mol. Biol. Evol. 25, 1307–1320 (July 2008).

16. Alley, E. C., Khimulya, G., Biswas, S., AlQuraishi, M. & Church, G. M. Unified rational protein engineering with sequence-based deep representation learning. en. Nat. Methods 16, 1315–1322 (Dec. 2019).

17. Lin, Z. et al. Evolutionary-scale prediction of atomic-level protein structure with a language model. Science 379, 1123–1130 (2023).

18. Madani, A. et al. Large language models generate functional protein sequences across diverse families. en. Nat. Biotechnol. 41, 1099–1106 (Jan. 2023).

19. Alamdari, S. et al. Protein generation with evolutionary diffusion: sequence is all you need. bioRxiv, 2023.09.11.556673 (Sept. 2023).

20. Ding, F. & Steinhardt, J. Protein language models are biased by unequal sequence sampling across the tree of life. bioRxiv (Mar. 2024).

21. Hou, C., Liu, D., Zafar, A. & Shen, Y. Understanding language model scaling on protein fitness prediction. en. bioRxivorg, 2025.04.25.650688 (Aug. 2025).

22. Jagota, M. et al. Cross-protein transfer learning substantially improves disease variant prediction. en. Genome Biol. 24, 182 (Aug. 2023).

23. Felsenstein, J. Maximum likelihood and minimum-steps methods for estimating evolutionary trees from data on discrete characters. Syst. Biol. 22, 240–249 (Sept. 1973).

24. Prillo, S. et al. CherryML: scalable maximum likelihood estimation of phylogenetic models. en. Nat. Methods 20, 1232–1236 (Aug. 2023).

25. Tenthorey, J. L. et al. Indels allow antiviral proteins to evolve functional novelty inaccessible by missense mutations. en. bioRxiv, 2024.05.07.592993 (May 2024).

26. Vaswani, A. et al. Attention is All you Need in Advances in Neural Information Processing Systems (eds Guyon, I. et al.) 30 (Curran Associates, Inc., 2017).

27. Sutskever, I., Vinyals, O. & Le, Q. V. Sequence to sequence learning with Neural Networks. arXiv [cs.CL] (Sept. 2014).

28. Rives, A. et al. Biological structure and function emerge from scaling unsupervised learning to 250 million protein sequences. Proceedings of the National Academy of Sciences 118, e2016239118 (2021).

29. Rao, R., Meier, J., Sercu, T., Ovchinnikov, S. & Rives, A. Transformer protein language models are unsupervised structure learners. en. Biorxiv, 2020.12.15.422761 (Dec. 2020).

30. Devlin, J., Chang, M.-W., Lee, K. & Toutanova, K. BERT: Pre-training of deep bidirectional Transformers for language understanding. arXiv [cs.CL] (Oct. 2018).

31. Yang, J. et al. Improved protein structure prediction using predicted interresidue orientations. en. Proc. Natl. Acad. Sci. U. S. A. 117, 1496–1503 (Jan. 2020).

32. Zhang, Z. et al. Protein language models learn evolutionary statistics of interacting sequence motifs. en. Proc. Natl. Acad. Sci. U. S. A. 121, e2406285121 (Nov. 2024).

33. Roch, S. Toward extracting all phylogenetic information from matrices of evolutionary distances. en. Science 327, 1376–1379 (Mar. 2010).

34. Atteson, K. The performance of neighbor-joining methods of phylogenetic reconstruction. en. Algorithmica 25, 251–278 (June 1999).

35. Holtzman, A., Buys, J., Du, L., Forbes, M. & Choi, Y. The curious case of neural text degeneration. arXiv [cs.CL] (Apr. 2019).

36. Wu, R. et al. High-resolutionde novostructure prediction from primary sequence. bioRxiv, 2022.07.21.500999 (July 2022).

37. Fletcher, W. & Yang, Z. INDELible: a flexible simulator of biological sequence evolution. en. Mol. Biol. Evol. 26, 1879–1888 (Aug. 2009).

38. Trost, J. et al. Simulations of sequence evolution: How (Un)realistic they are and why. en. Mol. Biol. Evol. 41, msad277 (Jan. 2024).

39. Guindon, S. et al. New algorithms and methods to estimate maximum-likelihood phylogenies: assessing the performance of PhyML 3.0. en. Syst. Biol. 59, 307–321 (May 2010).

40. Höhler, D., Haag, J., Kozlov, A. M. & Stamatakis, A. A representative Performance Assessment of Maximum Likelihood based Phylogenetic Inference Tools. bioRxiv (Nov. 2022).

41. Westesson, O., Lunter, G., Paten, B. & Holmes, I. Accurate reconstruction of insertion-deletion histories by statistical phylogenetics. en. PLoS One 7, e34572 (Apr. 2012).

42. Strope, C. L., Scott, S. D. & Moriyama, E. N. indel-Seq-Gen: a new protein family simulator incorporating domains, motifs, and indels. en. Mol. Biol. Evol. 24, 640–649 (Mar. 2007).

43. Ly-Trong, N., Naser-Khdour, S., Lanfear, R. & Minh, B. Q. AliSim: A fast and versatile phylogenetic sequence simulator for the genomic era. en. Mol. Biol. Evol. 39, msac092 (May 2022).

44. Altschul, S. F., Gish, W., Miller, W., Myers, E. W. & Lipman, D. J. Basic local alignment search tool. en. J. Mol. Biol. 215, 403–410 (Oct. 1990).

45. Brohawn, S. G., del Mármol, J. & MacKinnon, R. Crystal structure of the human K2P TRAAK, a lipid- and mechano-sensitive K+ ion channel. en. Science 335, 436–441 (Jan. 2012).

46. Natale, A. M., Deal, P. E. & Minor Jr, D. L. Structural insights into the mechanisms and pharmacology of K2P potassium channels. en. J. Mol. Biol. 433, 166995 (Aug. 2021).

47. Van Kempen, M. et al. Fast and accurate protein structure search with Foldseek. en. Nat. Biotechnol. 42, 243–246 (Feb. 2024).

48. Holmes, I. H. Historian: accurate reconstruction of ancestral sequences and evolutionary rates. en. Bioinformatics 33, 1227–1229 (Apr. 2017).

49. Löytynoja, A. & Goldman, N. Phylogeny-aware gap placement prevents errors in sequence alignment and evolutionary analysis. en. Science 320, 1632–1635 (June 2008).

50. Roney, J. P. & Ovchinnikov, S. State-of-the-art estimation of protein model accuracy using AlphaFold. en. Phys. Rev. Lett. 129, 238101 (Dec. 2022).

51. Jumper, J. et al. Highly accurate protein structure prediction with AlphaFold. en. Nature 596, 583–589 (July 2021).

52. Desmarais, J. J. et al. DABs are inorganic carbon pumps found throughout prokaryotic phyla. en. Nat Microbiol 4, 2204–2215 (Dec. 2019).

53. Merlin, C., Masters, M., McAteer, S. & Coulson, A. Why is carbonic anhydrase essential to Escherichia coli? en. J. Bacteriol. 185, 6415–6424 (Nov. 2003).

54. Kimber, M. S. & Pai, E. F. The active site architecture of Pisum sativum beta-carbonic anhydrase is a mirror image of that of alpha-carbonic anhydrases. en. EMBO J. 19, 1407–1418 (Apr. 2000).

55. Rowlett, R. S. et al. Kinetic characterization of wild-type and proton transfer-impaired variants of beta-carbonic anhydrase from Arabidopsis thaliana. en. Arch. Biochem. Biophys. 404, 197–209 (Aug. 2002).

56. Smith, K. S., Ingram-Smith, C. & Ferry, J. G. Roles of the conserved aspartate and arginine in the catalytic mechanism of an archaeal beta-class carbonic anhydrase. en. J. Bacteriol. 184, 4240–4245 (Aug. 2002).

57. Bracey, M. H., Christiansen, J., Tovar, P., Cramer, S. P. & Bartlett, S. G. Spinach carbonic anhydrase: investigation of the zinc-binding ligands by site-directed mutagenesis, elemental analysis, and EXAFS. en. Biochemistry 33, 13126–13131 (Nov. 1994).

58. Khalifah, R. G. The carbon dioxide hydration activity of carbonic anhydrase. I. Stop-flow kinetic studies on the native human isoenzymes B and C. en. J. Biol. Chem. 246, 2561–2573 (Apr. 1971).

59. Bar-Even, A. et al. The moderately efficient enzyme: evolutionary and physicochemical trends shaping enzyme parameters. en. Biochemistry 50, 4402–4410 (May 2011).

60. Hopf, T. A. et al. Mutation effects predicted from sequence co-variation. Nature biotechnology 35, 128–135 (2017).

61. Riesselman, A. J., Ingraham, J. B. & Marks, D. S. Deep generative models of genetic variation capture the effects of mutations. Nature methods 15, 816–822 (2018).

62. Meier, J. et al. Language models enable zero-shot prediction of the effects of mutations on protein function. Advances in neural information processing systems 34, 29287–29303 (2021).

63. Nijkamp, E., Ruffolo, J. A., Weinstein, E. N., Naik, N. & Madani, A. Progen2: exploring the boundaries of protein language models. Cell systems 14, 968–978 (2023).

64. Gordon, C. W., Lu, A. X. & Abbeel, P. Protein Language Model Fitness is a Matter of Preference in The Thirteenth International Conference on Learning Representations (2025).

65. Pugh, C. W., Nuñez-Valencia, P. G., Dias, M. & Frazer, J. From Likelihood to Fitness: Improving Variant Effect Prediction in Protein and Genome Language Models. bioRxiv, 2025–05 (2025).

66. Notin, P. et al. Proteingym: Large-scale benchmarks for protein fitness prediction and design. Advances in Neural Information Processing Systems 36, 64331–64379 (2023).

67. Prillo, S., Wu, W. Y. & Song, Y. S. Ultrafast classical phylogenetic method beats large protein language models on variant effect prediction in The Thirty-eighth Annual Conference on Neural Information Processing Systems (2024).

68. Vig, J. et al. BERTology Meets Biology: Interpreting Attention in Protein Language Models. en. Cold Spring Harbor Laboratory, 2020.06.26.174417 (June 2020).

69. Trinquier, J., Uguzzoni, G., Pagnani, A., Zamponi, F. & Weigt, M. Efficient generative modeling of protein sequences using simple autoregressive models. en. Nat. Commun. 12, 1–11 (Oct. 2021).

70. Mistry, J. et al. Pfam: The protein families database in 2021. Nucleic Acids Research 49, D412–D419 (2021).

71. Needleman, S. B. & Wunsch, C. D. A general method applicable to the search for similarities in the amino acid sequence of two proteins. en. J. Mol. Biol. 48, 443–453 (Mar. 1970).

72. Nesterenko, L., Blassel, L., Veber, P., Boussau, B. & Jacob, L. Phyloformer: Fast, accurate, and versatile phylogenetic reconstruction with deep neural networks. en. Mol. Biol. Evol. 42 (Apr. 2025).

73. Xie, V. C., Pu, J., Metzger, B. P., Thornton, J. W. & Dickinson, B. C. Contingency and chance erase necessity in the experimental evolution of ancestral proteins. en. Elife 10 (June 2021).

74. Yang, K. K., Wu, Z. & Arnold, F. H. Machine-learning-guided directed evolution for protein engineering. en. Nat. Methods 16, 687–694 (Aug. 2019).

75. Paszke, A. et al. PyTorch: An imperative style, high-performance deep learning library. arXiv [cs.LG] (Dec. 2019).

76. Su, J. et al. RoFormer: Enhanced Transformer with Rotary Position Embedding. arXiv [cs.CL] (Apr. 2021).

77. Sahoo, S. S. et al. Simple and effective masked diffusion language models. arXiv [cs.CL] (June 2024).

78. Nichol, A. et al. GLIDE: Towards photorealistic image generation and editing with text-guided diffusion models. arXiv [cs.CV] (Dec. 2021).

79. Williams, R. J. & Zipser, D. A learning algorithm for continually running fully recurrent neural networks. en. Neural Comput. 1, 270–280 (June 1989).

80. Dao, T., Fu, D. Y., Ermon, S., Rudra, A. & Ré, C. FlashAttention: Fast and Memory-Efficient Exact Attention with IO-Awareness. arXiv [cs.LG] (May 2022).

81. Katoh, K., Misawa, K., Kuma, K.-I. & Miyata, T. MAFFT: a novel method for rapid multiple sequence alignment based on fast Fourier transform. en. Nucleic Acids Res. 30, 3059–3066 (July 2002).

82. Heinzinger, M. et al. Bilingual language model for protein sequence and structure. en. NAR Genom. Bioinform. 6, lqae150 (Dec. 2024).

83. Kim, G. et al. Easy and accurate protein structure prediction using ColabFold. en. Nat. Protoc. 20, 620–642 (Mar. 2025).

84. Llinares-López, F., Berthet, Q., Blondel, M., Teboul, O. & Vert, J.-P. Deep embedding and alignment of protein sequences. en. Nat. Methods 20, 104–111 (Jan. 2023).

85. Loshchilov, I. & Hutter, F. Decoupled weight decay regularization. arXiv preprint arXiv:1711.05101 (2017).

86. Liljas, A. & Laurberg, M. A wheel invented three times. The molecular structures of the three carbonic anhydrases. en. EMBO Rep. 1, 16–17 (July 2000).

