## Supplementary Materials for "Deep models of protein evolution in time generate realistic evolutionary trajectories and functional proteins"

**This PDF file includes:**

Materials and Methods
Supplementary Text
Supplementary Table S1
Supplementary Figures S1 to S7

### Materials and Methods

#### Dataset Creation

The training and validation dataset was created as in [24]. Briefly, the TrRosetta dataset [31] was downloaded from its repository (https://github.com/gjoni/trRosetta). This dataset contains 15,051 multiple sequence alignments (MSAs) in a3m format. This format retains all alignment information - deletions relative to the query sequence are represented as gap characters ('–'), whereas insertions are retained as lower-case characters. Switching between a standard alignment and a set of full length sequences (not aligned) is thus easy.

The 15,051 MSAs were subsampled to at most 2048 sequences each, and randomly split into 14,498 train and 553 test (fully held-out) sets. From each MSA, a tree was estimated using FastTree [3] using the WAG [14] rate matrix (1 rate category). These trees were split at an early branching point to yield approximately balance subtrees. One tree was used in model training, the other was reserved as an in-family held-out set to test generalization within protein families seen in training. Cherries were picked greedily, starting from the closest pair of sequences in each subtree. For the classical (WAG and LG) models, aligned sequences were used. For PEINT training and validation, the original sequences, with gaps removed and insertions relative to query retained were used.

#### Model Design and Implementation

The PEINT transformer was implemented in Pytorch [75]. Our final model consists of 5 transformer encoder layers (on top of the 30 provided by ESM2-150M [frozen]), as well as 5 standard Transformer decoder layers with 20 attention heads and an embedding dimension of 640, with 57M trainable parameters. The encoder layers use the self-attention operator to build progressively more sophisticated representation of the input sequence. The decoder layers alternate between masked self-attention, that learns from already decoded tokens, and cross attention to the encoder representation at a similar layer. Concretely, rather than having each layer in the decoder cross attend to the final hidden representation learned by the encoder, decoder layer 1 cross attends to the hidden representation from encoder layer 1, and so on. The model's initial embedding layer, as well as its final projection layer (that is weight-tied to the embedding layer) are also imported from ESM2-150M and are frozen. Positional encoding is accomplished using rotary positional encoding (RoPe) [76] to maintain consistency with ESM2, which also uses RoPe, as well as for the flexibility it offers for sequences of different lengths.

The time encoding was inspired by the diffusion model literature [77, 78], in which continuous time values are provided as conditioning tokens to a neural network that is designed to denoise an input. Concretely, given time $t$, and a set of predefined geometrically space frequencies $\text{freqs} = \{f_0, \ldots, f_{d//2}\}$ where $d$ is the model embedding dimension, we generate a time embedding as $\text{emb}(t) = (\cos(\text{freqs} * t)^\intercal, \sin(\text{freqs} * t)^\intercal)^\intercal$. This time vector is added to the input sequence $y$ after the embedding layer, similar to how fixed positional encodings are provided to transformer models [26].

The starting sequence $x$ was provided to the encoder with a prepended special 'start' token (`cls`) and appended 'end' token (`eos`), following the convention from ESM2. We randomly selected 15% of tokens to replace with a special `mask` token for masked language model training on the encoder. The final sequence $y$ was input with the same 'start' token (`cls`) to right shift the sequence by one token. The objective function on the decoder is a standard autoregressive loss (next token prediction) using teacher forced maximum likelihood training [79]. We also added the 'end' token to the input $y$ so PEINT would learn to terminate sequences during generation. The PEINT model was trained with a composite loss function that was equal parts masked language modeling (encoder) and autoregressive (decoder). We implemented our model using FlashAttention [80] and trained on 2× A100 GPUs (NVIDIA) for ∼24 to 36 hours (with early stopping imposed when validation loss stopped decreasing), with a batch size of roughly 250 thousand tokens, and a learning rate of $4 \times 10^{-4}$ with a weight decay of 0.01 applied to non-bias parameters. To maintain consistency with the sequence lengths that ESM2 was trained on, we only trained PEINT on sequences of less than 1022 amino acids.

The WAG and LG models were fit to the same data as the PEINT model to allow for a direct comparison. The LG model allowed for 4 discrete site rates. Models were fit using the same composite likelihood objective, using the CherryML package[24].

##### Evaluating Transition Likelihoods

Evaluating transition likelihoods with both classical and deep models of protein evolution is made challenging by their fundamental asymmetry - classical models require alignments, while deep models work with non-aligned sequences. To enable a direct comparison between these model classes, we implemented a position masking strategy that accounts for both insertions and deletions in an alignment. Starting from an `a3m`-formatted alignment, where insertions relative to query are denoted by lowercase letters, we constructed (1) an alignment mask that identifies aligned (uppercase) positions in the sequences, and (2) a gap mask that indexes the positions of gap characters in the alignment. We could then evaluate transition likelihoods on the cherry pairs $(x, y, t)$ in this alignment. For classical models, we evaluate the transition likelihood per alignment site $i$ in a cherry as usual: $l_i = \exp(Qt\alpha_i)[x_i, y_i]$, where $Q$ is the transition rate matrix of the model in question (LG or WAG), and $\alpha_i$ is the site rate at position $i$ ($\alpha_i$ is uniformly 1 for WAG) estimated by FastTree, and $t$ is the time separating the sequences $x$ and $y$. For the PEINT model, we evaluated per-site transition likelihoods across full-length (insertions retained) unaligned sequences (gap characters removed), producing predictions for all residues in $y$ including insertions. We could then apply the two masks sequentially to map these positions to the classical model's aligned coordinate system. The alignment mask retains aligned positions in the alignment (discarding insertions), and the gap mask identifies deletion sites in the aligned sequences. We assigned identical likelihood values to positions containing a gap in the aligned sequences as a means of normalizing out their impact. This approach allowed us to ensure that the transition likelihoods reflect model performance on residue substitutions at homologous positions while avoiding artifacts from differential treatment of insertions and deletions. We evaluated transition likelihoods on 512 cherries from each of the roughly 550 held-out test trees, as well as 256 cherries from 550 randomly sampled in-family held-out subtrees.

##### Simulating Evolution on Single Branches

In order to efficiently generate candidate sequences ($y$) given a starting sequence $x$ and branch length ($t$), we implemented a cached version of the PEINT model that minimizes repeat computation of static intermediates. After the first token has been generated, we omit the encoder completely; its contribution to the decoding process occurs through$k, v$ vectors in the cross attention operation that can be stored in memory, rather than recomputed. We found this encoder caching led to a ∼5 fold increase in generation speed. We additionally cache the self-attention $k, v$ vectors as each token in $y$ is generated, as is standard in decoder-only, GPT style models. In total, these caching mechanisms improved generation speeds by ∼10 fold. Simulation was initiated by providing the model with a `cls` token, and generation proceeded autoregressively until the model output an `eos` token. To avoid generating non-amino acid tokens that are present in the ESM vocabulary, we set their logit scores to $-\inf$ prior to the softmax operation and sampling from the output probabilities. Tokens were sampled from these output distributions without further modification, as various probability truncation strategies resulted in significantly reduced diversity across all branch lengths. To simulate single branches using classical models, we first generated transition probability matrices for that branch length, where $P(t) = \exp(Qt)$. For the LG model, we used the site rates estimated by FastTree for a particular MSA, and generated site-rate specific transition probability matrices $P_{WAG}(T) = \exp(Qt\alpha_i)$ where $\alpha_i$ indicates the site rate at position $i$. We then randomly sampled each site independently.

##### Branch Length Estimation

To estimate branch lengths for pairs of sequences, we perform stochastic gradient descent on the PEINT negative log likelihood for those sequences as a function of time. We use the Adam optimizer with an initial learning rate of $1 \times 10^{-1}$ that decreases exponentially with $\gamma = 0.99$. We found that estimates typically converged in fewer than 80 steps.

##### Simulating Evolution on Trees

We selected trees from the held-out set of TrRosetta MSAs that did not contain sequences longer than 1022 amino acids. For each tree, we selected the leaf node with the median sequence length to be the new root, and re-rooted the tree on this leaf node. The subtree that did not contain this new leaf node was designated as the evaluation set. Sequences from this subtree serve as an internal control for the expected diversity of sequences at this distance from the root. To simulate evolution along trees using PEINT, we use a breadth-first traversal strategy implemented using a queue-based data structure to efficiently utilize GPU parallelization. This queue-based approach enables batching of multiple sequence generation tasks across different tree branches. In early experiments, we found that PEINT would occasionally generate repetitive sequence stretches, that would grow progressively longer with subsequent generation cycles. These repeat expansions usually indicated that the `eos` was not generated at a reasonable place in the output sequence ($y$). To mitigate this rare behavior, we employed a length based filtering scheme in which sequences outside a specific ratio of parent length are rejected (typically, we allowed 10% expansion or contraction of the parent length). For each batch iteration, we generate $N$ (here, 4) candidate sequences at once, and randomly select from the sequences that pass our filter. If no sequences pass filtering criteria, the parent node is re-queued with an incremented retry counter (maximum 3 attempts). Upon exceeding retry limits, a random sequence from the generated candidates is selected to ensure tree completion. Once all the leaves have been generated, the simulation algorithm stops. For downstream structural and conservation analyses, sequences simulated using PEINT were aligned to the empirical MSA using MAFFT[81], with the `--keeplength` flag.

For classical models, we used the AliSim [43] simulation package, using either the WAG or LG model with 4 rate categories (`LG+G4`) as the evolutionary model, as these are the most commonly used models in the field. The core AliSim command used to generate simulated MSAs was:

```
IQTree2 --alisim <output> -te <tree> -s <msa> --root-seq <msa,root_seq> -o
↪  <root_id> --out-format fasta -seed 42 --write-all
```

with the addendum `-m WAG` if using the WAG model, or `-m LG+G4 --site-rate MODEL` if using LG with 4 site rates. Here, the `-te <tree>` and `-s <msa>` correspond to the rerooted tree from the PEINT simulation, and the empirical MSA from TrRosetta, respectively. Importantly, when providing the original MSA as an input to the simulator, the empirical gap pattern is maintained, leading to an output MSA with the same gap register as the real sequences. This obviates the need to re-align the sequences, and ensures that alignment errors are minimized.

**Simulation Benchmarking**

For the evolutionary simulations, we selected $\sim 550$ trees from the held-out set and re-rooted them on the leaf node with median sequence length. The tree was then split into two subtrees: a root-proximal subtree and an evaluation subtree (the other subtree not containing the root node). The evaluation subtree served as an internal control for expected sequence divergence at equivalent evolutionary distances.

Mutational Burden In Simulations To assess the realism of substitution rates along evolutionary trajectories, we quantified the mutation burden accumulated across branches in simulated versus natural sequence evolution. We restricted this analysis to the evaluation subtrees as described above. For natural sequences, we first inferred ancestral sequences using Historian [48] with phylogeny-aware gap placement, as described in further detail below. The reconstructed ancestor of the evaluation subtree was then used as input to AliSim to simulate evolution down the evaluation subtree topology using either the WAG or LG+G4 model, following the simulation parameters described in the Simulating Evolution on Trees section. For PEINT simulations, we performed simulations over the entire re-rooted tree and subsequently restricted our analysis to the evaluation subtree. Since re-rooting can cause node renaming, we matched corresponding branches between simulated and natural trees by identifying branches with identical sets of descendant leaf nodes. For each matched branch, we calculated the number of amino acid substitutions that occurred between the parent and child nodes. We excluded indel events from this analysis to avoid potential biases introduced by alignment errors. Substitution counts were then aggregated across all branches and models, allowing us to examine the relationship between substitution rates and branch length. This enabled direct comparison of the mutation accumulation patterns between PEINT, classical evolutionary models (WAG and LG+G4), and natural sequence evolution.

Conservation Analysis To evaluate whether simulated sequences maintain biologically relevant conservation patterns, we performed a conservation analysis on both amino acid sequences and 3Di structural states. Using the simulation subtree alignment, we identified highly conserved sites, defined as positions where a single amino acid state occurred in $> 70\%$ of sequences. For PEINT-simulated sequences, leaf nodes were aligned to the empirical MSA using MAFFT [81] with the `--keeplength` flag. For classical model simulations (WAG and LG+G4), sequences were already aligned due to AliSim's gap pattern preservation, obviating the need for re-alignment.

At each conserved site, we quantified the divergence between simulated and natural amino acid distributions using the Jensen-Shannon Divergence (JSD):

$$\text{JSD}(P \parallel Q) = \frac{1}{2} D_{KL}(P \parallel M) + \frac{1}{2} D_{KL}(Q \parallel M)$$

Where $M$ is a mixture distribution ($M = \frac{1}{2}(P+Q)$) and $D_{KL}$ is the Kullback-Leibler divergence. Here, P represents the amino acid frequency distribution at a conserved site in the simulated sequences, and Q represents the distribution in the evaluation subtree. To establish a baseline for expected divergence, we also calculated the JSD between real sequences from the root-proximal and evaluation subtrees.

For 3Di state conservation analysis, we first converted protein sequences to 3Di structural alphabet states using the ProstT5 model [82], then followed the identical procedure described above to identify conserved 3Di positions and calculate divergences between simulated and real leaf sequences.

Historian To compare indel event rates between natural and PEINT-simulated sequence evolution, we used the Historian [48] package. Historian uses a phylogeny aware gap-placement algorithm that has been shown to lead to less biased inference of insertions and deletions compared to standard sequence alignment algorithms. This allows us to reconstruct evolutionary trajectories from a set of real sequences and their associated inferred tree. For the PEINT simulations, we could provide the ancestral states to Historian, as those are generated in the process of decoding, granting us direct access to these sequences. We restricted our analysis to the evaluation subtree (Figure 3a). As input to Historian, we provided the subtree (`tree_file`) and a guide multiple sequence alignment (`msa_file`) generated using MAFFT. This guide alignment is treated as a "hint" and we adjusted the band parameter (that allows for sliding of gaps in the guide alignment) to allow for substantial refinement of the alignment. The actual command used to infer substitution and indel events was:

```
historian recon -guide <msa_file> -tree, <tree_file> -ancseq -output fasta
↪  -allspan -band 40 -refine
```

We could then record evolutionary events along the tree.

Structural Prediction We used a variety of methods to evaluate the structural validity of sequences generated using PEINT or classical models of protein evolution. These include two methods that predict structures from single sequences (OmegaFold, AF2Rank). For these approaches, we randomly selected 30 leaves from the evaluation subtree of each simulation tree and report the median pLDDT across these leaves. For OmegaFold, the sequences were provided as a single fasta file. For AF2Rank, which uses single sequences combined with a structural template for prediction, we first aligned the leaf sequences to the TrRosetta exemplar structure of that protein family, omitting insertions relative to the structure. For AlphaFold2, we provided the MSAs directly for structure prediction. We first aligned the outputs of the various simulations to the sequence of the exemplar structure from TrRosetta for each family. We then provided this multiple sequence alignment as an input to AlphaFold2 through local ColabFold [83].

##### Model Interpretation

We adapted the categorical Jacobian formulation from [32] to be used with PEINT for model interpretation. PEINT is an autoregressive model which learns conditional likelihoods of the form: $p(Y|X,t) = \prod_{i=1}^{L} p(Y_i|Y_{<i}, X, t)$. Using a similar notation to [32], we will use $Y(y_i \to a_k)$ to denote the sequence obtained by mutating position $i$ of sequence $Y$ into the $k$th amino acid, $a_k$. The categorical Jacobian of the model, $J$, is a tensor of shape $L \times A \times L \times A$, where $L$ is the length of sequence and $A$ is the size of the alphabet. For the decoder categorical Jacobian used for contact prediction, we set $X = Y = s_{wt}$ where $s_{wt}$ is the wild-type sequence with a known structure, then computed each entry as:

$$J_{ikjl} = \log p\big[Y_j = a_l | Y(y_i \to a_k)_{<j}, X, t\big] - \log p\big[Y_j = a_l | Y_{<j}, X, t\big]$$

Due to the autoregressive nature of the decoder, we set all entries where $i > j$ to zeroes. In other words, for each position $j$ in the outputs, we mutate each position $i$ in the inputs into every possible amino acid, then collect the changes in the output logits for position $j$ over all amino acids. To obtain a $L \times L$ coupling matrix to be used for contact prediction, following [32], we mean centered and symmetrized $J$, then took the Frobenius norm over the two amino acid dimensions and applied the average product correction (APC). We compared contact prediction using PEINT's decoder categorical Jacobian with ESM2's categorical Jacobian on 553 sequences in the test set families.

For the encoder categorical Jacobian used pairwise alignment between sequence $X$ and $Y$, we used a similar calculation without the autoregressive constraints:

$$J_{ikjl} = \log p\big[Y_j = a_l | Y_{<j}, X(x_i \to a_k), t\big] - \log p\big[Y_j = a_l | Y_{<j}, X, t\big]$$

To evaluate the accuracy of the encoder-decoder categorical Jacobian in terms of pairwise alignment prediction, we used a setup similar to that of [84]. In brief, we downloaded curated Pfam-A [70] seed alignments for 1,000 protein domains. For each domain, we randomly selected a pair of aligned sequences from the seed MSA. This pairwise alignment was treated as the ground truth. We then used the un-aligned versions of the pair of sequences (gaps removed) as inputs to either a classical aligner (implemented using Biotite using the Blosum62 matrix) or to the PEINT categorical Jacobian. We converted the ground truth and classical aligner-inferred alignments match-state trace matrices. The PEINT categorical Jacobian output was symmetrized, row-softmaxed, and finally binarized using a threshold of 0.2. To assay the quality of classical, or learned alignments from the PEINT categorical Jacobian, we calculate an F1 score ($F1 = 2 * \frac{\text{precision}*\text{recall}}{\text{precision}+\text{recall}}$ over match positions (with gap placement being the complement of matches). We stratified alignment performance by the empirical number of differences between the sequences in the Pfam reference alignment.

##### Variant Effect Prediction

To train a PEINT model to be used for variant effect prediction on the ProteinGym substitution benchmark [66], we first extracted transitions for each of the families using the MSAs provided by the ProteinGym benchmark. As we only trained the model on sequences shorter than 1,022 amino acids in length to maintain consistency with ESM2, we imposed the same restriction at evaluation time, and we report our performance relative to other models on the 201 of the 217 total families that fit this criterion. We followed the same procedure as for the TrRosetta dataset to infer evolutionary trees and extract evolutionary transitions from these MSAs. The model shared very similar architecture and training hyperparameters to the PEINT model trained on the TrRosetta data, with the difference that it only has 2 encoder and 2 decoder layers. We used the AdamW optimizer [85] with a learning rate of $2 \times 10^{-4}$, weight decay of 0.01, batch size of 32, and gradient accumulation of 6 batches. The model was trained on a single A100 GPU for ∼2 hours with early stopping imposed based on the Spearman correlation with the DMS measurements on 25 randomly sampled families.

For evaluation, we score mutant via $score(x^{\text{mut}}, x^{\text{wt}}) = \log p(x^{\text{mut}} | x^{\text{wt}}, t)$, where $t$ is a hyper-parameter controlling that size of the evolutionary neighborhood around the wild-type. We used $t = 1$ for variant effect prediction [67]. We followed the convention of the ProteinGym benchmark and computed the Spearman correlation for each family, then report the Spearman correlation aggregated over all families for each assay type as the primary metric. For ESM2, we used the scores reported by the ProteinGym benchmark.

##### *In-vivo* characterization of Carbonic Anhydrases

Genes coding the Pea $\beta$-carbonic anhydrase ($\beta$CA) (PDB ID: 1EKJ), the Pea $\beta\text{CA}^{D152N}$, and simulated variants were ordered as gene blocks (Twist Bioscience). All genes were cloned into a pLTet vector ($\text{Kan}^R$) that places the genes under control of a Tet-on promoter. Plasmids were transformed into CAFree *Escherichia coli* [52], and grown overnight at 37°C in permissive (5%) $CO_2$ conditions in Luria Broth (LB) media with $50\frac{\mu g}{\text{mL}}$Kanamycin. Stationary phase cultures were diluted $1 : 100$ into LB Media + Kanamycin, and grown at permissive (5%) $CO_2$ until mid log-phase ($OD_{600}$ ∼0.3). Colonies were then diluted to an $OD_{600}$ of 0.001 into fresh LB + Kan, and supplemented with

200nM Anhydrotetracycline (ATC) ((4S,4aS,12aS)-4-(Dimethylamino)-1,4,4a,5,12,12a-hexahydro-3,10,11,12a-tetrahydroxy-6-methyl-1,12-dioxo-2-naphthacenecarboxamide) to induce expression. Growth curves were measured on a Tecan Spark (Tecan) microplate reader, at a non-growth-permissive $CO_2$ concentration of 1% with orbital shaking at 37∘C for 72 hours using a humidity casette to ensure wells did not dry out. Wells that showed growth were diluted into fresh LB with $50\frac{\mu g}{\text{mL}}$Kanamycin, grown overnight at 37°C and sequenced to ensure that growth arose as a result of the correct gene.

##### ***In-vitro* characterization of Carbonic Anhydrases**

Carbonic anhydrase genes were cloned into a pET-SUMO vector; the resultant protein has an amino terminal 14×His-SUMO tag. Proteins were expressed in LOBSTR (KeraFast) *Escherichia coli* cells. Two liters of luria broth (L.B.) media were inoculated to 1% final volume with an overnight culture of transformed cells. Cells were grown at 37°C until they reached an $OD_{600}$ of ∼0.6 and protein expression was induced by addition of 1mM IPTG overnight at 20°C. Cells were harvested by centrifugation, lysed in lysis buffer (500mM NaCl, 20mM HEPES pH 7.5, 10$\mu$M $Zn^{2+}$, 5mM $\beta$-mercaptoethanol) using sonication, and the soluble fraction was harvested by centrifugation at 20,000×g. Soluble lysate was supplemented with 20mM Imidazole, and loaded onto 3mL of Ni-NTA resin (Qiagen). Resin was washed with 50 column volumes high salt wash buffer (500mM NaCl, 20mM HEPES pH 7.5, 10$\mu$M $Zn^{2+}$, 5mM $\beta$-mercaptoethanol, 40mM Imidazole), 15 column volumes low salt wash buffer (250mM NaCl, 20mM HEPES pH 7.5, 10$\mu$M $Zn^{2+}$, 5mM $\beta$-mercaptoethanol, 40mM Imidazole), and eluted in low salt wash buffer supplemented with 250mM Imidazole. Eluted protein was dialyzed overnight into low salt wash buffer without imidazole, in the presence of 1:100 SUMO Protease. Residual uncleaved protein, His-SUMO tag, and SUMO protease were removed by passing the dialyzed eluant over ∼1mL Ni-NTA resin and collecting the flowthrough, which was then concentrated and purified using size exclusion chromatography (SEC) on either a Superose 6 or Superdex 200 column in low salt wash buffer (without imidazole). Peak fractions were assayed by SDS-PAGE and pooled, concentrated to $> 50\mu$M, and flash frozen in liquid nitrogen. The final, purified protein likely contained a mix of different oligomeric states [86].

*In vitro* $CO_2$ hydration rates were measured using the method of Khalifa [58] on a Kintek AutoSF-120 (Kintek Corp) stopped flow device. One syringe was filled with a saturated solution of $CO_2$ in water (roughly ∼34mM $CO_2$), formed by bubbling $CO_2$ gas into milli-Q water for $> 30$ minutes. The other syringe contained 100mM MOPS pH 7.5 (with ionic strength adjusted using $Na_2SO_4$), 100$\mu$M p-nitrophenol, and either 2$\mu$M (Seq1360, Pea$\beta$CA$^{D152N}$ or 200nM (Pea$\beta$CA$^{WT}$) protein, or nothing (uncatalyzed reaction). Reactions were initialized by rapid mixing of equal amounts of the contents of both syringes using the stopped flow device, and reaction progress was monitored by measuring absorbance at 478nm. Each curve shown represents the average of 4 independent reaction measurements (individual curves shown in Supplementary Figure S5).

### Supplementary Text

#### General Primer on Evolutionary Modeling

Formally, a model of protein evolution is a conditional probability distribution $p(y|x,t)$ which, given a starting protein sequence $x$ and an elapsed time $t$, describes the distribution for the evolved sequence $y$. Classically, protein evolution has been modeled with continuous-time Markov chains (CTMCs) which 1) require sequences to be aligned with gap characters ('–') so that they have the same length, and 2) assume that each site of a protein evolves independently such that the joint conditional probability $p(y|x,t)$ factorizes over sites $i$ as $\prod_i p(y_i|x_i,t)$.

Unfortunately, we rarely get to observe samples $(x,y,t)$ from an evolutionary model $p(y|x,t)$, since only present-day proteins are observed. Instead, we obtain observations from a related model, called a *phylogenetic model*, which is constructed out of the evolutionary model of interest. The construction is as follows: The phylogenetic model is a tree-structured Bayesian network with tree structure parameterized by $\mathcal{T}$ and conditional probability distributions given by $p(y|x,t)$. The root distribution of the tree is given by a prior probability $\pi_{\text{root}}(x)$, typically given by the stationary distribution. The phylogenetic model is a statistical model for the dataset $D$ of protein sequences at the leaves of the tree $\mathcal{T}$, generated as follows:

1. First, the protein sequence $x$ for the root of the tree $\mathcal{T}$ is drawn from $\pi_{\text{root}}(x)$.

2. Having sampled the protein sequence $x$ for internal node $u$ of the tree, for the edge $(u,v,t)$ of the tree, we sample protein sequence $y$ for node $v$ following the conditional distribution $p(y|x,t)$. This captures protein evolution forward in time.

3. The process continues until protein sequences are sampled for all leaves, yielding $D$.

Crucially, this is a latent variable model in which the protein sequences at internal nodes of the tree $\mathcal{T}$ are unobserved, complicating statistical inference of the underlying model of $p(y|x,t)$.

In general, one has access to many trees $\mathcal{T}_1, \mathcal{T}_2, \ldots, \mathcal{T}_m$ each corresponding to one of $m$ different protein families (e.g. , insulin, hemoglobin, etc.) and datasets of observed leaf proteins $D_1, D_2, \ldots, D_m$ for these respective trees. The dataset $D_i$ of leaf sequences is traditionally arranged into a multiple sequence alignment (MSA), which makes all sequences have the same length by adding gap characters (denoted with '–'). It is assumed that the datasets $D_i$ are independent and each follows the above phylogenetic model parameterized by $\pi_{\text{root}}(x)$, $p(y|x,t)$ and the tree $\mathcal{T}$. If the evolutionary model is family-specific, then this is captured by a more general contextual model $p(y|x,t,c)$ where $c$ contains information about the protein family (e.g. , the 3D structure of the protein). The goal then becomes to estimate the model of protein evolution $p(y|x,t)$ from the leaf observations $D_1, \ldots, D_m$ and the trees $\mathcal{T}_1, \ldots, \mathcal{T}_m$. In what follows, let $n$ be the typical number of sequences per MSA and $l$ the typical protein sequence length. Classical datasets such as Pfam have $m \approx 10,000, n \approx 1,000, l \approx 200$, for a total on the order of $10^9$ residues (i.e., tokens).

Historically, models of protein evolution have relied on the independent-sites assumption and on CTMCs. The WAG model [14] posits that each site in a protein evolves independently according to a $20 \times 20$ rate matrix $Q$, normalized such that the expected number of substitutions in a unit of time is equal to 1. In this model, protein evolution is defined by $p(y|x,t) = \prod_{i=1}^{l} \exp(tQ)[x_i, y_i]$, where $\exp(tQ)[x_i, y_i]$ denotes the $[x_i, y_i]$ element of the matrix exponential $\exp(tQ)$, and $x$ and $y$ are aligned protein sequences of length $l$; gaps are treated as ignorable missing data. The most popular model used to date is the LG model [15], which allows for site-rate variation via family-specific site-rates $c = (\alpha_1, \alpha_2, \ldots, \alpha_l)$, yielding $p(y|x,t,c) = \prod_{i=1}^{l} \exp(\alpha_i tQ)[x_i, y_i]$.

Despite the simplicity of classical models, learning classical models of protein evolution is not an easy task. Generally, maximum likelihood estimation of phylogenetic models requires dealing with the unobserved ancestral states, for which expectation-maximization (EM) is the method of choice. For independent-site models such as the WAG model of protein evolution, the data log-likelihood can be computed with dynamic programming [23] (DP) in time $\Omega(mnls^2)$ where $s = 20$ is the number of amino acids, and more generally, the number of states of the CTMC. This comes from having to recursively compute the quantities $\text{dp}[v][a]$, which encode the (log-)probability that running the phylogenetic model starting at node $v$ in state $a$ produces the observed data in the leaves descending from $v$, as in the classical forward-backward algorithm for inference in classical hidden Markov models. Unfortunately, if the independent-sites assumption is dropped, then exact computation of the full likelihood of the data becomes intractable since one needs to integrate out over exponentially many ancestral proteins. In other words, one would need to compute $\text{dp}[v][a]$ where $a$ no longer takes just $s = 20$ values, but rather $s = 20^l$ values where $l$ is the protein sequence length.

CherryML [24] proposes replacing the full joint likelihood of the data by a composite likelihood over cherries in the trees. Following the notation in [24], letting $c_i$ be the number of cherries in tree $\mathcal{T}_i$, and letting $(u^i_j, v^i_j)_{1\le j\le c_i}$ be the cherries in tree $\mathcal{T}_i$, CherryML builds the composite (log-)likelihood:

$$l_{\text{comp}} = \sum_{i=1}^{m}\sum_{j=1}^{c_i} \log p_{\text{phylo}}(D_i[u^i_j] | D_i[v^i_j], \mathcal{T}_i) \tag{S1}$$

Here $D_i[u^i_j]$ indexes the protein sequence in dataset $D_i$ corresponding to the leaf node $u^i_j$ of the tree. Since the composite likelihood involves only pairs of sequences at a time, it is much easier to compute than the full joint likelihood. In fact, if one assumes the model of protein evolution is parameterized by a time-reversible CTMC then the term $p(D_i[u^i_j] | D_i[v^i_j], \mathcal{T}_i)$ depends only on the distance between $u^i_j$ and $v^i_j$ in tree $\mathcal{T}_i$. Thus, denoting by $t^i_j$ the pairwise distance between $u^i_j$ and $v^i_j$ in tree $\mathcal{T}_i$, the composite likelihood can be rewritten in terms of the likelihood $p$ of the underlying model of protein evolution as:

$$l_{\text{comp-rev}} = \sum_{i=1}^{m}\sum_{j=1}^{c_i} \log p(D_i[u^i_j] | D_i[v^i_j], t^i_j), \tag{S2}$$

where $c_i$ denotes the number of cherries $(u^i_j, v^i_j)$ in tree $\mathcal{T}_i$, $t^i_j$ the pairwise distance between $u^i_j$ and $v^i_j$ in tree $\mathcal{T}_i$, and $D_i[u^i_j]$ indexes the sequence in MSA $D_i$ corresponding to the leaf node $u^i_j$ of $\mathcal{T}_i$. It was shown that this massively speeds up MLE, several orders of magnitude faster than previous methods, while retaining consistency and high statistical efficiency [24].

We do not constrain our deep model of protein evolution to be time-reversible. Instead, we consider an unconstrained deep parameterization of $p(y|x, t)$. We do this for a number of reasons. For one, constraining the model to be time-reversible is non-trivial outside of classical independent-sites CTMCs. Also, the premise of deep learning is that with reasonably flexible architectures and enough training data, any interesting structure in the data will be captured by the model without hardcoding this structure ourselves. In other words, we let the data speak for itself. If the true process of protein evolution is indeed time-reversible, then the unconstrained deep model may learn this structure without imposing these constraints explicitly. The biggest caveat is that when the model of protein evolution is not time-reversible, then the reversible composite likelihood $l_{\text{comp-rev}}$ of (S2) no longer guarantees consistent estimation of model parameters. Nonetheless, (S2) provides an approximate means to learn general deep models of protein evolution. In light of this, it is more appropriate to think of the loss in (S2) as supervising the model to predict, for a given starting sequence $x$ and an evolutionary time $t$, the distribution of evolutionary '*neighbors*' of $x$ at evolutionary distance $t$ rather than only direct evolutionary '*descendants*' of $x$; these two notions agree for time-reversible models. Such a model is clearly still useful since it will capture the evolutionary landscape around a given protein, as needed for phylogenetic tree reconstruction, variant effect prediction, and broadly for protein design applications. Thus, our work represents a tradeoff between expressive power of the model, and consistency of the estimation procedure: while classical models are simplistic and can be consistently estimated, deep models are highly expressive but we resort to a non-consistent estimation procedure.

### Supplementary Table

**Table S1**: **Model Hyperparameters.** Peint model hyperparamters

| Hyperparameter | Value |
|---|---|
| Pretrained Encoder | ESM2 (150M) |
| Encoder layers | 5 |
| Decoder layers | 5 |
| Embedding Dimension | 640 |
| Batch size (tokens) | $\sim$250,000 |
| Learning Rate | $4 \times 10^{-4}$ |
| Weight decay | 0.01 |
| Trainable Parameters | $\sim$57 M |

#### Supplementary Figures

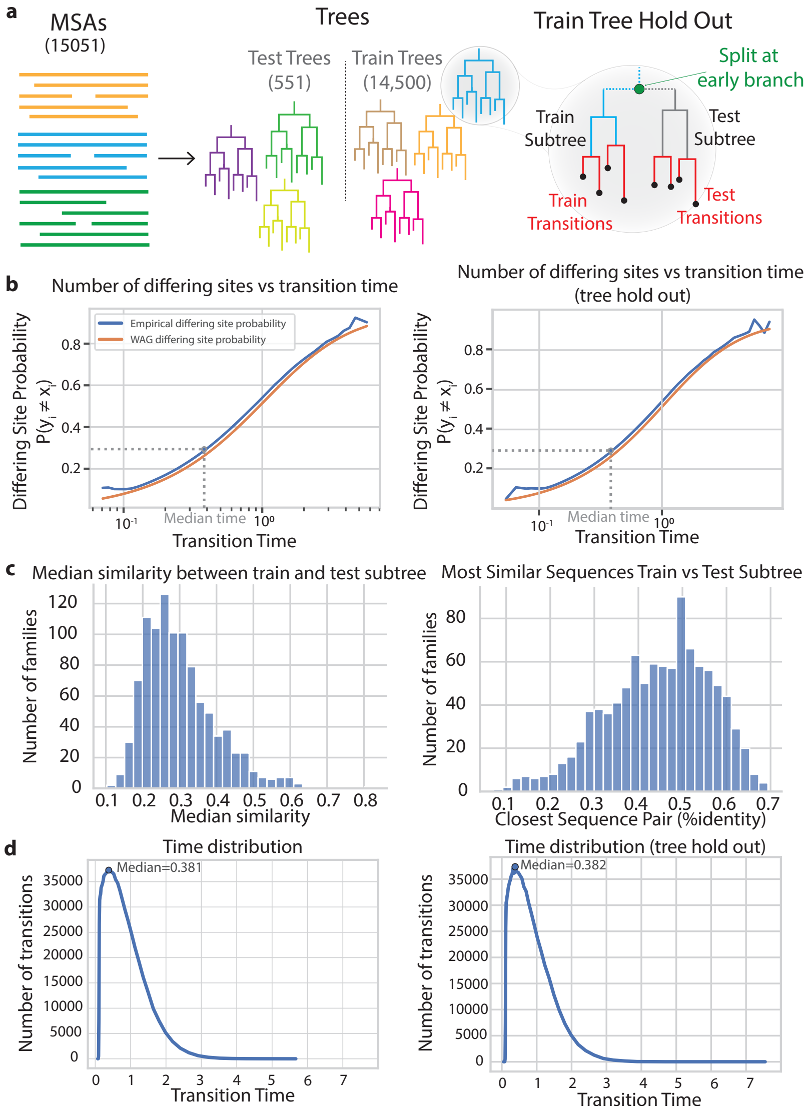


**Figure S1**: **TrRosetta Dataset Statistics. a)** Dataset partitioning - 500 trees are held out as the test set. The remaining 14,500 trees are split into two equally-sized subtrees, one is included in the training set, the other is held out. **b)** Empirical and theoretical measures of sequence differences as a function of time. Orange curve indicates theoretical mutational load, blue indicates empirical observations. The median transition has $\sim$30% sites that differ between $x$ and $y$. Left is from the training subtrees, right is from the held-out subtrees, showing similar statistics. **c)** Median all-vs-all pairwise percent identity (left) and closest pair of sequences (right) between held-out and training subtrees. The two trees show minimal overlap in sequence space. **d)** Distribution of transition times from the cherries in the training and held-out subtree sets. Training and held-out subtrees show similar distributions of cherry branch lengths.

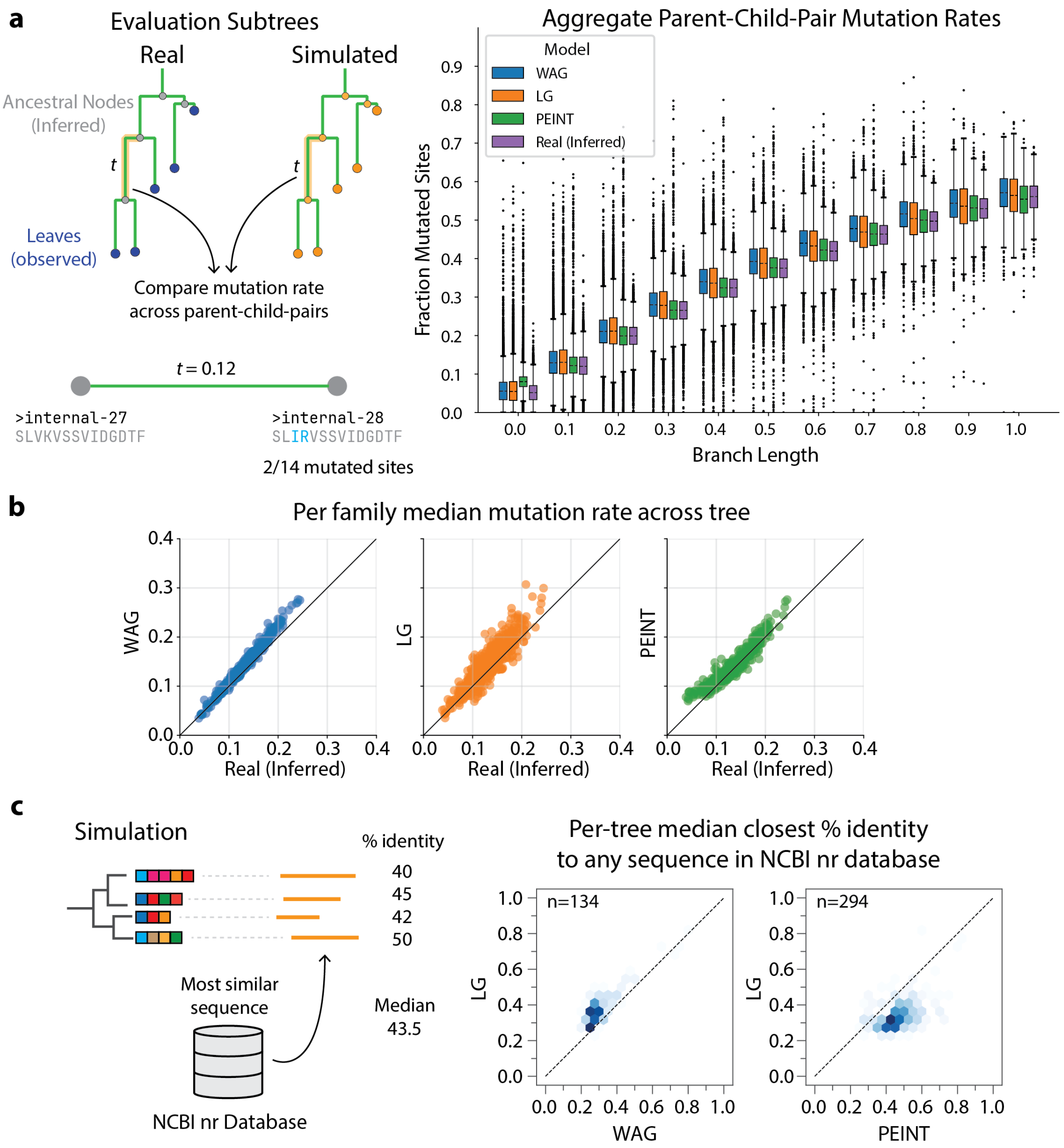


**Figure S2**: **Simulations generate sequences with natural level of mutations a)** Mutation rates of various models across parent-child pairs were compared to those inferred using ancestral reconstructions from real trees and aggregated by branch length. **b)** Same data setup as in **a**, but the median mutation rate for each tree was taken over all parent-child pairs. The subtrees all contained identical branches. **c)** Density plot showing similarity of simulated sequences to any extant sequence in the BLAST nonredundant protein database (nr). For each family, the closest sequence (by % identity) to each leaf node in the nr database was identified, and the median closest distance was calculated. Comparison between different simulation methods show they all generate sequences that are distinct from extant sequences (< 50% sequence identity) WAG and LG simulations frequently produced nonsensical sequences that yielded no hits in BLAST, and only trees for which at least 10 sequences had a match in BLAST were included, leading to different numbers of families for the different analyses.

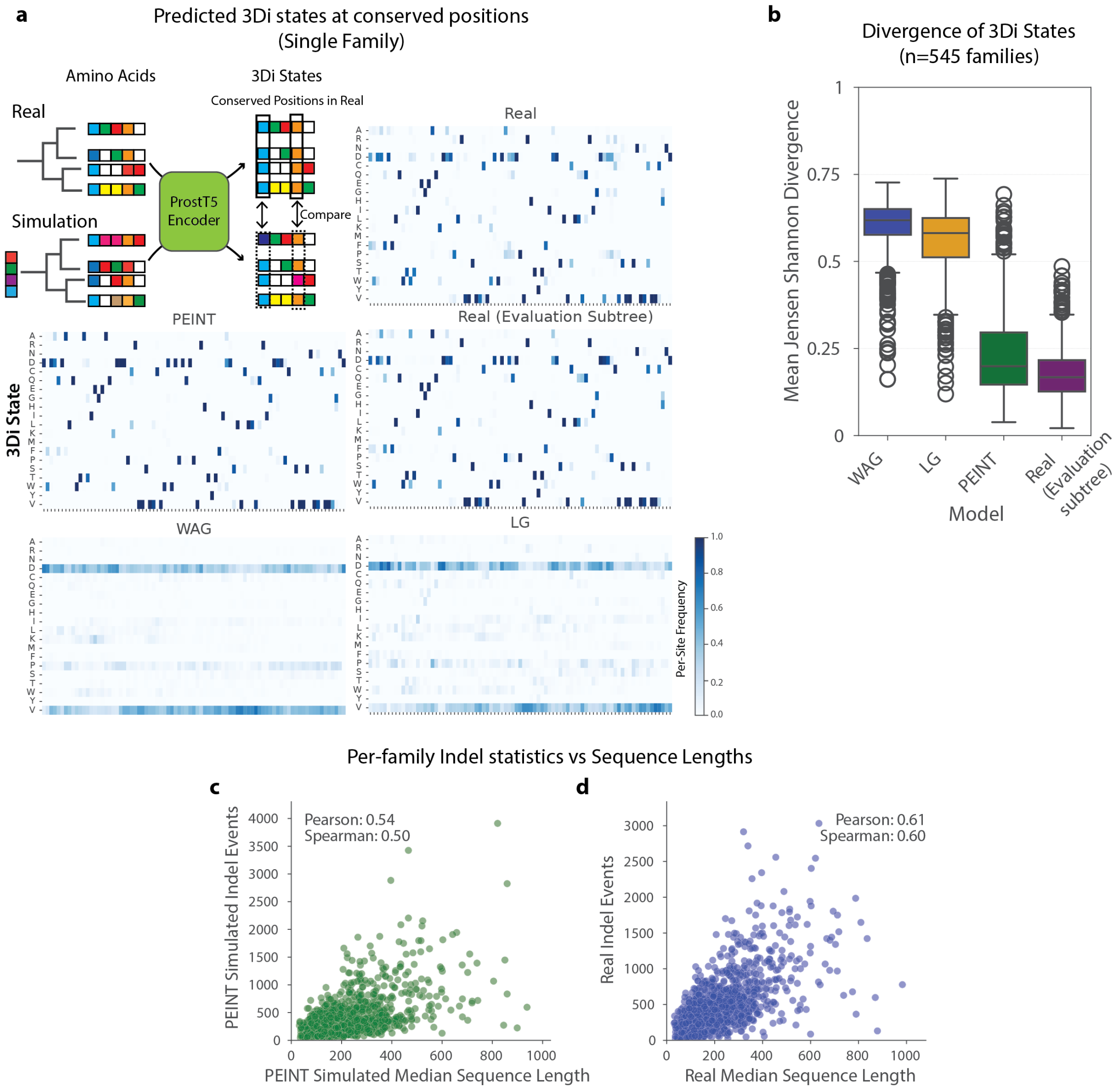


**Figure S3**: **PEINT simulates realistic evolution. a)**: 3Di conservation analysis schematic. For each protein family, real and simulated sequences were translated to 3Di states using the ProstT5 model. Conserved sites were identified from the real MSA, and usage patterns of 3di states at those sites were compared between real sequences and simulated ones. Matrices show per-site 3di state frequencies in both subtrees of real sequences, as well as the various simulation methods. **b)**: Aggregate mean per-site Jensen-Shannon divergence between real and simulated 3Di frequencies at empirically conserved sites across all simulation families (n=545). **c)**: Comparison of the number of per-family indel events in PEINT simulated sequences as a function of median sequence length in that family. **d)**: Comparison of the number of per-family indel events inreal sequences as a function of median sequence length in that family.

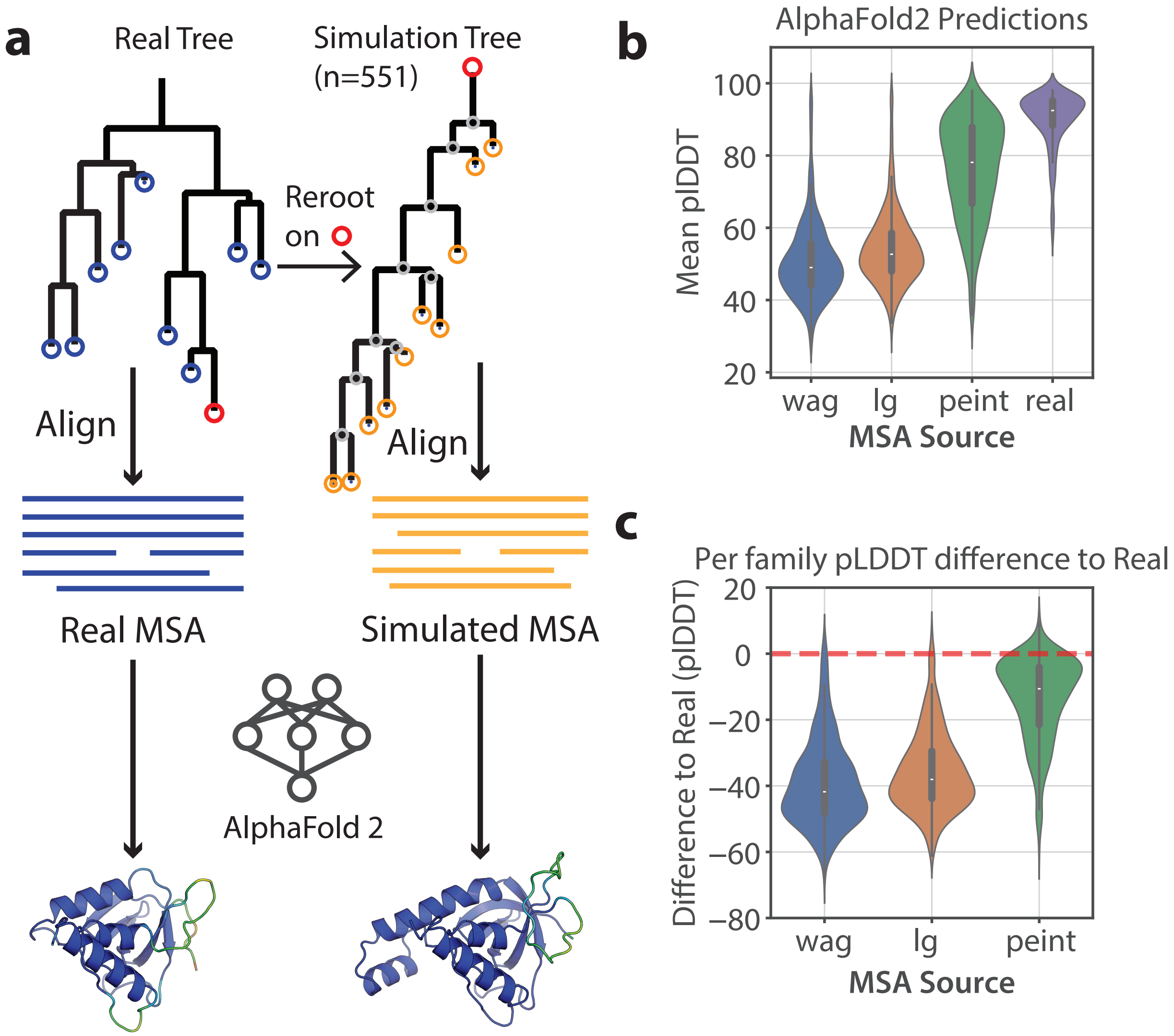


**Figure S4**: **AlphaFold2 readily folds structures from PEINT Simulations. a)**: For each protein family, we compare the quality of simulated sequences to real sequences using AlphaFold2 as a discriminator. The simulated sequences are aligned and provided to AlphaFold2 as the multiple sequence alignment to aid in structure prediction. AlphaFold2 pLDDT per family shown (**(b)**, as well as per-family pLDDT difference to the structure predicted using a real MSA (**(c)**. Classical simulators (WAG, LG) generate poor MSAs for the purpose of structure prediction. PEINT (green) generates MSAs that are similar in quality to those composed of real sequences (purple).

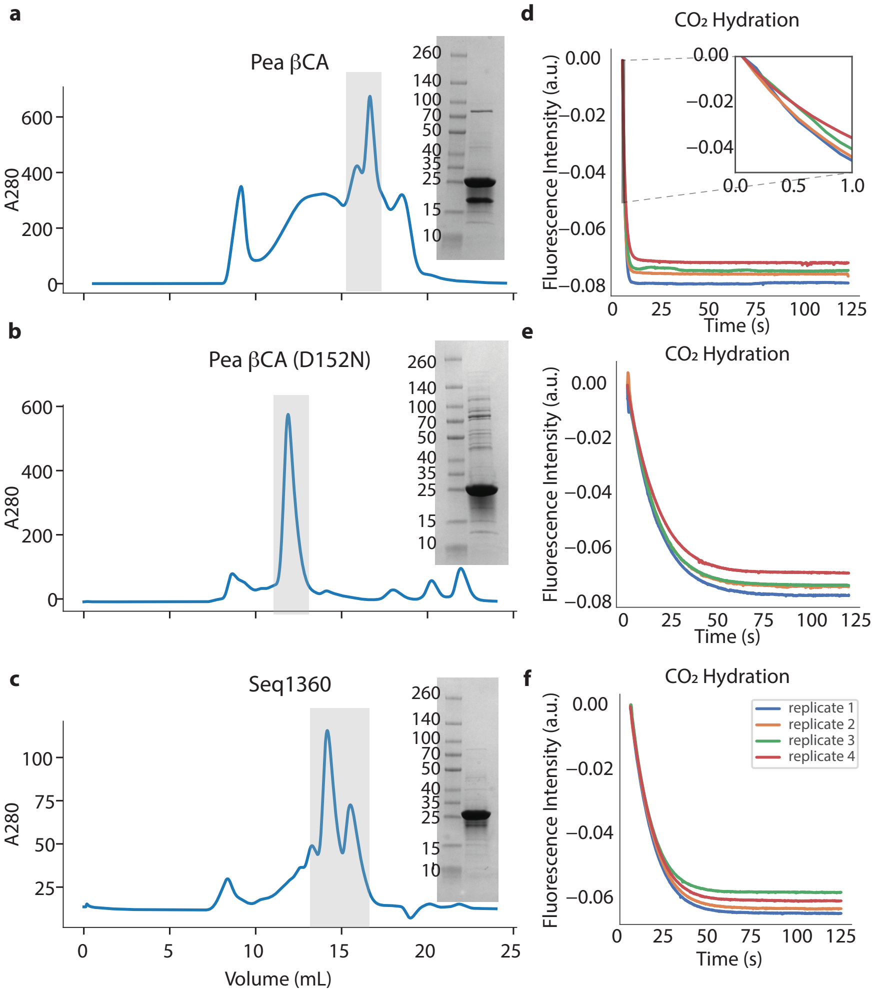


**Figure S5**: **Purification and biochemical characterization of carbonic anhydrases.** Overview of purification and stopped-flow traces for the $\beta$-carbonic anhydrases used in the study. **a-c**: size exclusion chromatograms, pooled fractions (gray) and SDS-PAGE gel of the final protein product. **d-f**: stopped flow traces of the various proteins at 17mM $CO_2$, replicates shown in different colors.

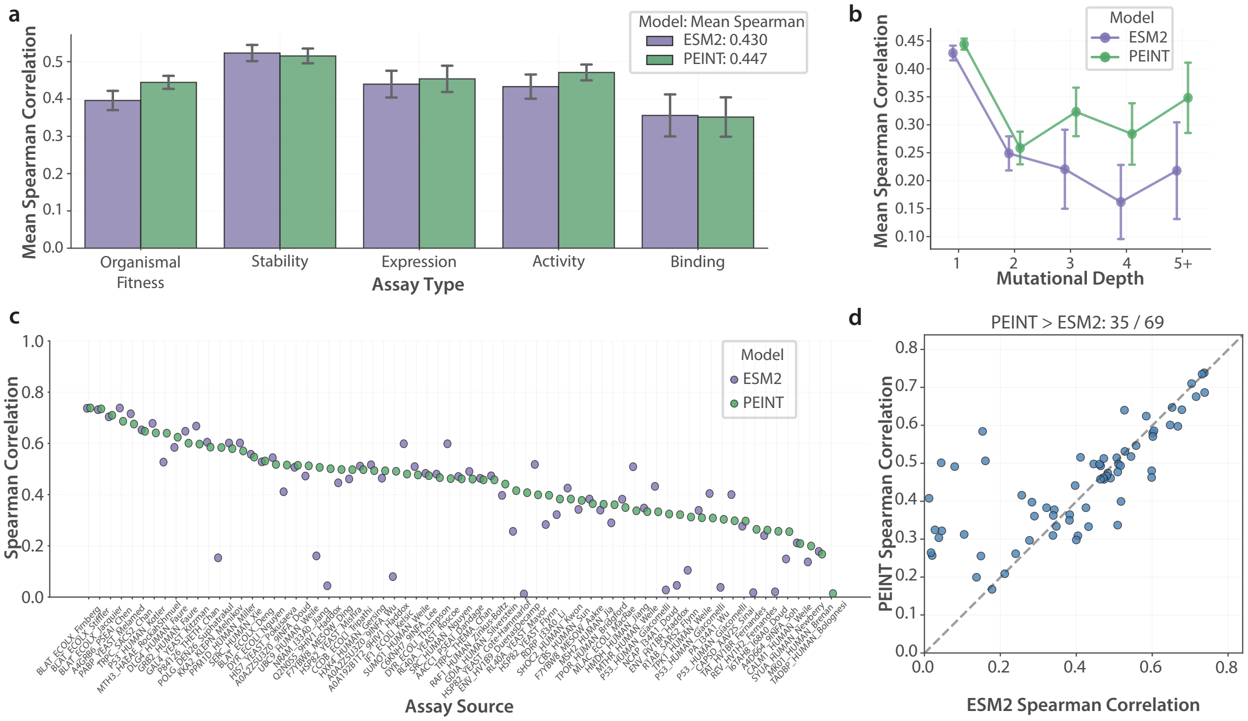


**Figure S6**: **PEINT improves variant effect prediction on a larger ESM model. a)** Mean Spearman correlation on 201 families in the ProteinGym substitution benchmark [66]. The families are stratified by assay type. The height of each bar represents the mean of the per-family Spearman correlations for each assay type. Error bars represent the standard deviation across proteins within each assay type. For each model, the average over the mean Spearman correlations for the five assay types is shown in the figure legend. "ESM2" is the 650M parameters ESM2 model, and "PEINT" is a PEINT model trained on transitions of the DMS families, using the frozen 650M parameters ESM2 in its encoder. **b)** Mean Spearman correlation stratified by mutational depths. **c)** Spearman correlation for each family in the ProteinGym substitution benchmark with the assay type "Organismal Fitness", where the assays measure the extent to which mutations affect an organism's growth rate. Each protein family is denoted by its identifier in the ProteinGym dataset. **d)** ESM2 vs. PEINT Spearman correlation for each family in the ProteinGym substitution benchmark with the assay type "Organismal Fitness".

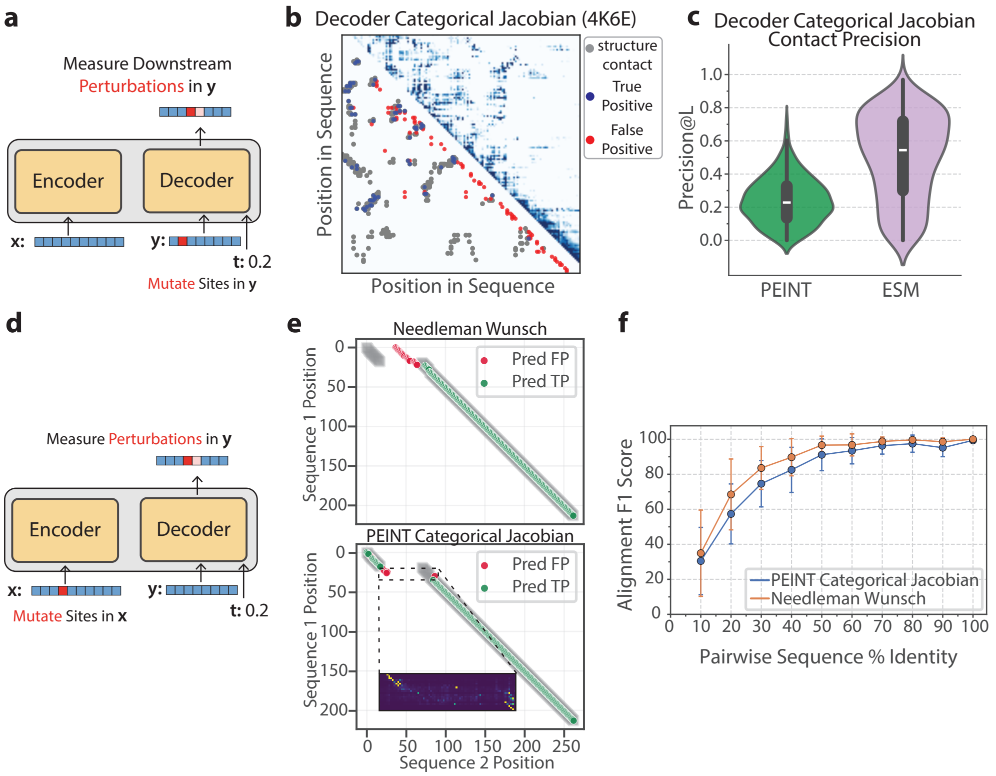


**Figure S7**: **PEINT Interpretability.** Top: Categorical Jacobian [32] analysis of the decoder half of the PEINT model. **a)** Cartoon showing the categorical Jacobian approach for the decoder. The $x$ sequence is fixed, and each position in $y$'s input are mutated, and perturbations are measured at downstream positions as a logit ratio to non-mutated. **b)** Example output (top right) categorical Jacobian matrix, overlayed with the contact map (bottom left) from the structure of the input sequence (PDB ID: 4K6E). The top $L$ strongest categorical Jacobian couplings are colored blue if they correspond to a pair of residues in contact with one another, and red if not. **c)** Aggregated contact precision of the Top $L$ categorical Jacobian couplings across 553 sequences from test-set families (green) compared to ESM (encoder-only) categorical jacobian couplings (purple). Bottom: Categorical Jacobian analysis *between* PEINT encoder and decoder blocks. **d)** Cartoon showing the categorical Jacobian calculation for the decoder. The $x$ sequence is mutated one site at a time, and perturbations to each position in the output of $y$ are measured. **e)** Example for a single pair of sequences from a Pfam seed alignment. The ground truth alignment from Pfam is shown as gray squares. **Top**: The trace matrix from a classical pairwise alignment algorithm (Needleman-Wunsch) is shown as circles, colored if each aligned site is correct (green) or incorrect (red). **Bottom**: PEINT's encoder-decoder categorical Jacobian matrix is shown, with couplings colored green if they correspond to correctly aligned residue pairs, and red if not. Inset: the raw values from the categorical Jacobian are shown at a stretch of inserted residues suggesting a 'soft' alignment in the gap region. **f)** Aggregate metrics as in **e** are shown over 1000 randomly selected Pfam domain seed alignments. Predicted alignment F1 scores are shown as a function of the pairwise sequence identity from the ground truth Pfam-seed alignment. Information flow between PEINT's encoder and decoder suggests a learned alignment.
